# Elucidating the regulation of glucose tolerance through the interaction between the reaction product and active site pocket residues of a β-glucosidase from *Halothermothrix orenii*

**DOI:** 10.1101/844506

**Authors:** Sushant K Sinha, Shibashis Das, Sukanya Konar, Pradip Kr. Ghorai, Rahul Das, Supratim Datta

## Abstract

β-glucosidase catalyzes the hydrolysis of β-1,4 linkage between two glucose molecules in cello-oligosaccharides and is prone to inhibition by the reaction product glucose. Relieving the glucose inhibition of β-glucosidase is a significant challenge. Towards the goal of understanding how glucose interacts with β-glucosidase, we expressed in *Escherichia coli*, the Hore_15280 gene encoding a β-glucosidase in *Halothermothrix orenii*. Our results show that the enzyme is glucose tolerant, and its activity stimulated in the presence of up to 0.5 M glucose. NMR analyses show the unexpected interactions between glucose and the β-glucosidase at lower concentrations of glucose that however does not lead to enzyme inhibition. We identified non-conserved residues at the aglycone-binding and the gatekeeper site and show that increased hydrophobicity at the pocket entrance and a reduction in steric hindrances are critical towards enhanced substrate accessibility and significant improvement in activity. Analysis of structures and in combination with molecular dynamics simulations show that glucose increases the accessibility of the substrate by enhancing the structural flexibility of the active site pocket and may explain the stimulation in specific activity up to 0.5 M glucose. Such novel regulation of β-glucosidase activity by its reaction product may offer novel ways of engineering glucose tolerance.

## 1. Introduction

Microbes express the enzymes required for the conversion of polysaccharides in lignocellulosic biomass to produce sugars. Through a fermentative process, the sugars can then be converted to biofuels by the same or other microbes. The enzymes that break down the polysaccharides into fermentable sugars are collectively known as cellulase. The minimum set of required enzymes in this cellulase mix (cellobiohydrolase, endoglucanase, and β-glucosidase) work synergistically to deconstruct the biomass [1-3]. Endoglucanase (EC 3.2.1.4) randomly cleave the β-1,4 glycosidic linkages of cellulose; cellobiohydrolase (EC 3.2.1.91 and 3.2.1.176) attack the cellulose chain ends to produce cellobiose (a dimer of glucose linked by a β-1,4 glycosidic bond); and β-glucosidase (EC 3.2.1.21) hydrolyze cellobiose into two molecules of glucose. The inhibition of cellobiose hydrolysis by β-glucosidase reaction product glucose is recognized as the limiting step in the conversion of lignocellulosic biomass to sugars [4]. The separate hydrolysis and fermentation (SHF) methodology, a commonly used biofuel production strategy is prone to product inhibition. Another efficient and economic cellulose hydrolysis setup under high-gravity fermentation also requires a high biomass loading and enzymes tolerant to the molar concentrations of glucose produced during the reaction [5]. Relieving glucose inhibition would result in the rapid increase in hydrolysis activity by the β-glucosidase and more economical biomass hydrolysis [6]. Relieving this product inhibition is thus a significant challenge.

The inhibition constant (*K*_i,app_) of glucose on the chromogenic model β-glucosidase substrate, *p*-nitrophenyl D-glucopyranoside (*p*NPGlc), spans many orders of magnitude with a few naturally occurring β-glucosidase with *K*_i, app_ in the molar range [7-12]. The glucose-induced inhibition has been attributed to a broader active site pocket entrance that facilitates increased glucose access to the enzyme active site [13]. Put another way, glucose tolerance was proposed to be a consequence of a narrower and deeper active site pocket that impedes access to the active site [13]. These observations do not, however, explain the ability of the enzyme pocket to distinguish between the reaction product glucose and the substrate cellobiose. Our sequence comparisons with other highly glucose tolerant β-glucosidase such as O08324, A0A0F7KKB7, and Q8T0W7 suggest that many of the residues previously implicated for glucose tolerance are non-conserved and therefore those specific residues may not play a role in glucose tolerance [14-16].

Therefore, we have embarked on a program to understand the role of the active site pocket in glucose tolerance of β-glucosidase and engineering glucose tolerance of low glucose tolerant enzymes. In this study, we used a β-glucosidase (B8CYA8) from the thermophilic and halophilic bacteria *Halothermothrix orenii* [17]. It was previously reported that B8CYA8 efficiently converts lactose to different transglycosylated products and hydrolyzes cellobiose to glucose [14, 18, 19]. Here we report that B8CYA8 is tolerant to high concentrations of glucose. Unexpectedly, we observed by NMR-based experiments that the enzyme interacts with glucose, even at low concentrations. Saturation-transfer difference (STD) NMR experiment verified that the glucose interaction with the B8CYA8 residues does not affect the enzyme. While the glucose may be expected to sterically hinder the access of substrate and inhibit B8CYA8 activity, the enzyme activity was stimulated by glucose. We identified conserved and non-conserved residues spanning the enzyme active site pocket that affects glucose tolerance to reveal the importance of amino acid residues across glycone, aglycone and gatekeeper sites of B8CYA8. A combination of beneficial mutants generated highly active variants of B8CYA8. Finally, based on our kinetic studies, structural analyses and molecular dynamics (MD) simulations, we propose a model to describe how glucose may regulate the stimulation and inhibition of the enzyme.

## 2. Methods

### 2.1. Chemicals

All chemicals used were of reagent grade. Restriction endonucleases, DNA ligase, and DNA polymerase were purchased from NEB (MA, USA). Primers were synthesized by Xceleris (India). All chromogenic substrates and chemicals were purchased from Sigma-Aldrich. The active fractions post-purification was pooled and concentrated using 30 kDa cut-off size membranes of Amicon-Ultra-15 (Merck Millipore, Bangalore, India). Plasmid purification and gel purification kits were obtained from Qiagen (Hilden, Germany).

### 2.2. Bacterial strains, culture conditions, and plasmids

The synthetic gene corresponding to the β-glucosidase from *H. orenii* was constructed (BankIt1930137BG_Halotherm KU867899) as reported previously and expressed in *Escherichia coli* Top 10F’ cells (Life Technologies, La Jolla, CA) [14]. The cells were centrifuged at 4000 ×*g* for 10 min at 4 °C and the pellet stored at −20 °C until purification of the protein.

### 2.3. Primer design, PCR, and cloning

All mutants were generated via a mega primer-based polymerase chain reaction (PCR) mutagenesis strategy [20]. Briefly, three primers were used - a template specific forward and reverse primer and the mutant primer either towards the forward or reverse direction, depending on the position of the mutation in the gene (Supplementary file, Table S7). The first PCR was run using the mutant primer and template-specific primer to generate the megaprimer containing the mutation. The megaprimer was extended during the second PCR by another sequence-specific primer. While the single mutants were generated from the wild-type DNA, the template containing single mutations were used to generate the double mutants. Primers were designed using OligoAnalyzer (IDT Technology) and ApE (ApE Plasmid Editor, version 2.0.49 by M. Wayne Davis). The DNA sequences encoding the mutants were obtained from both strands by automated DNA sequencing at the IISER Kolkata sequencing facility.

### 2.4. Protein expression and purification

B8CYA8 and mutants were purified using a protocol as detailed earlier, and purity was confirmed by 10 % SDS–PAGE [14]. The protein concentrations were determined by measuring the absorbance at 280 nm and using the extinction coefficient for respective enzyme variants calculated using the modified Edelhoch and Gill/Von Hippel methods on Expasy (http://web.expasy.org/protparam/).

### 2.5. Saturation transfer difference (STD) NMR of protein B8CYA8 and ligand glucose

The sample for NMR experiment was prepared in 20 mM potassium phosphate buffer, pH 7.0, and 99.9 % D_2_O, final B8CYA8 concentration was at 90 μM and glucose concentration at 20 mM. NMR spectra were recorded on a Bruker AV 500 spectrometer at 298 K. The saturation transfer difference (STD) spectra were recorded by setting the on- and off-resonance irradiation at −1 ppm and 30 ppm, respectively [21]. All spectra were recorded with 256 scans, four dummy scans, a spectral width of 8012 Hz, and 8 K points. The residual protein background signal was suppressed with the 30 ms T1ρ filter. In determining the ligand signal arising from direct saturation of ligand signals close to on-resonance pulse, a control sample with ligand only was used. All spectra were processed and analyzed with Topspin 3.5 (Bruker Biospin Corporation, MA, USA).

### 2.6. Enzyme activity assays

The pH dependence of B8CYA8 mutants was determined by measuring enzyme specific activities on *p*NPGlc in the pH range of 5.0 to 8.0 at 70 °C, after incubating the enzyme overnight at 4 °C in each buffer. The effect of temperature on enzyme activity for *p*NPGlc was measured between 55 to 80 °C while incubating in McIlvaine buffer, pH 6.5. Based on the initial rate measured, the amount of enzyme to be used, and the assay time was optimized. The specific activity of the mutants was assayed at the T_opt_ and pH_opt_ of each enzyme as per previously published protocol, using saturating concentrations of substrates, *p*NPGlc, and cellobiose (Clb) [22]. Clb hydrolysis produces two molecules of glucose and the calibration curve used was based on the glucose produced.

### 2.7. Kinetic Analysis of B8CYA8

The kinetic parameters of all the mutants were determined at various concentrations, ranging between 0.5 mM to 100 mM, of substrates *p*NPGlc and Clb as previously reported [14]. GraphPad PRISM version 7.0 (GraphPad Software, La Jolla, CA) was used to calculate all kinetic constants by a non-linear regression fit of the Michaelis-Menten equation.

### 2.8. Thermostability, half-life, residual specific activity assay, and T_m_

Enzymes were incubated in 100 mM HEPES buffer, pH 7.1 for wild-type enzyme, and McIlvaine buffer at pH_opt_ of each mutant at 70 °C. At regular time intervals, aliquots were taken out, centrifuged, and assayed for the residual specific activity. Half-life times were determined using the equation for a one-phase exponential decay in GraphPad PRISM. The residual specific activity of enzymes in the presence of glucose was determined at 70 °C upon 24 h incubation in the buffer containing 1 M glucose without substrate. Then the samples were cooled down, and specific residual activity was measured with 20 mM *p*NPGlc at the respective optimum conditions. For each sample, blanks without enzyme were subtracted for any background absorbance.

### 2.9. Measurement of synergy

The synergy of B8CYA8 and its mutants with commercial cellulase derived from *Trichoderma viride* (Sigma-Aldrich, St. Louis, USA) was measured on Avicel. The 200 μL of reaction contained 20 μg of cellulase, 3 μg of B8CYA8 or mutants, and 15 % (w/v) Avicel, in a buffer of pH 5.0. Sweet almond β-glucosidase (SRL, Chennai, India) was used as a control for B8CYA8. The reaction time course was followed until two hours at 37 °C. The reaction was terminated by heating at 95 °C for 10 min, and the glucose generated quantitated using a GOD-POD assay kit (Sigma-Aldrich, St. Louis, USA).

### 2.10. Molecular Dynamics Simulations

The X-ray crystallographic structure of the β-glucosidase was obtained from the protein databank (PDB: 4PTX)[18]. The energy minimized protein molecule (B8CYA8) was kept at the center of a cubic simulation box 120 Å long and then solvated with water. We used the TIP3P water model in all of our simulations [23]. Glucose molecules were added by using PACKMOL to obtain glucose concentrations [24]. All the potential parameters were obtained from the CHARMM36 force field. The simulations were performed by NAMD-2.9 simulation tools [25-27]. By taking initial configurations at different glucose concentrations, the conjugate gradient method was applied for 300000 steps to remove all energetically unfavorable contacts. The starting configurations were equilibrated for 3.0 ns in the NPT ensemble to fix the simulation box length. In NPT simulations, Noose-Hoover thermostat and barostat coupling constants were taken to be 0.5 ps and 2.0 ps, respectively [28, 29]. The pressure was kept constant at 1.0 atm. After the simulation box length was fixed, we equilibrated the system for 5.0 ns in the NVT ensemble. To analyze different system properties, a 50 ns production run was performed in the NVT ensemble. The chosen system temperature was kept constant by using the damping coefficient (γ) of 1.0 ps^−1^ by Langevin dynamics. Long-range interactions are handled by the particle mesh Ewald (PME) method with real space cut-off of 16 Å and 2 Å pair list cut-off [30-32]. We used 1-4 scaling factor in our simulations. The time step was 1.0 fs, and all the properties were computed from the trajectories stored at an interval of 4.0 ps during the production run.

## 3. Results

### 3.1. The effect of glucose on B8CYA8 specific activity

We assayed enzyme-specific activity in the presence of exogenously added glucose and observed that the presence of 0.5 M to 0.75 M glucose, stimulated B8CYA8 specific activity by 1.7-fold, unlike the typical enzymatic product inhibition profiles. Though further addition of glucose decreased B8CYA8 activity, around 125 % specific activity was retained in the presence of 1.5 M glucose (Fig. 1). Glucose has been commonly known to be a competitive inhibitor of β-glucosidase, wherein the apparent *K*_*m*_ increases with increasing glucose concentration without any change in *k*_*cat*_. However, in B8CYA8, both stimulation and inhibition are at play as both *K*_*m*_, and *k*_*cat*_ increases with an increase in glucose concentration (Supplementary file, Table S1).

**Figure 1.**
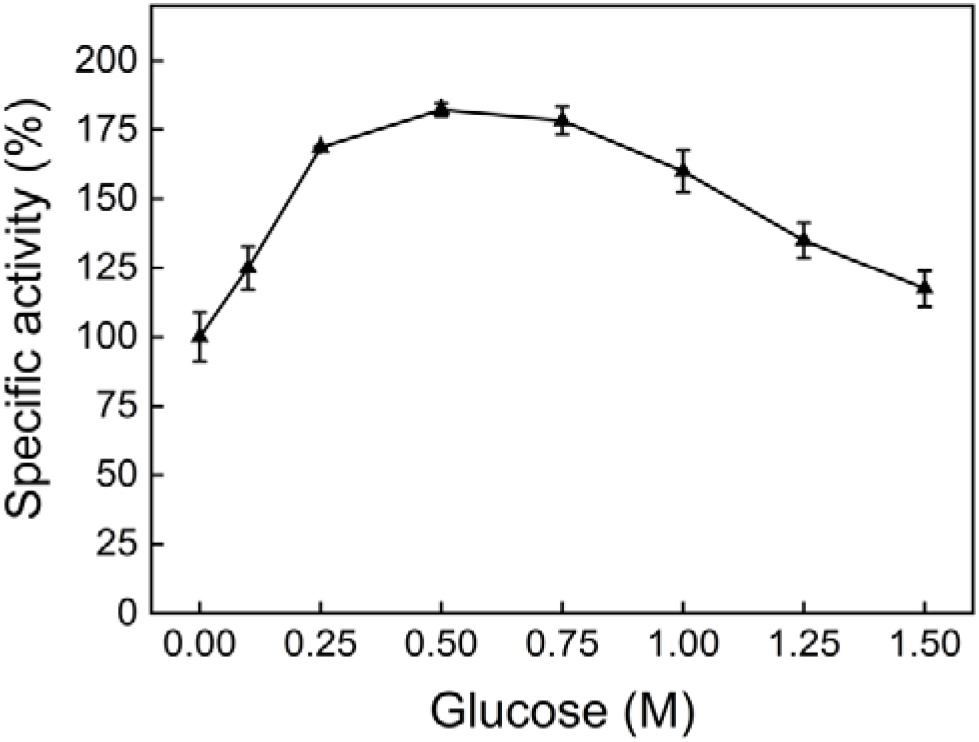
Effect of D-glucose on B8CYA8 specific activity using *p*NPGlc (20 mM) as substrate. The specific activity assay details are included in the materials and methods section.

The crystal structure of B8CYA8 complexed with glucose (PDB: 4PTX), show glucose trapped in the glycone binding region (−1 subsite) and occupying the substrate-binding site [18]. The site of glucose binding could indicate a competitive inhibition of B8CYA8 by glucose. Alternately, the presence of glucose could have been an artifact of the crystallization trials. Since the authors had soaked the crystal with a non-hydrolysable substrate, the glucose could have been trapped due to crystal packing. To ascertain if glucose indeed interacts with B8CYA8 in solution, we probed the glucose enzyme interaction by NMR.

### 3.2. Interaction of glucose and B8CYA8 by Saturation-transfer difference (STD) NMR experiment

The interaction of glucose with B8CYA8 was probed by 1D STD-based NMR experiments [21]. Fig. 2b shows the one-dimensional reference spectra of glucose, STD spectra of glucose alone as a control, and the STD spectra of glucose in the presence of B8CYA8. In the sample containing the protein and glucose, we could observe the 1D-NMR signal of the glucose when the on-resonance pulses were set at the aliphatic region of the protein, enabling the transfer of the magnetization to glucose (Fig 2b). In the control experiment, the STD spectra of only glucose did not produce any signal for glucose. The transfer of magnetization to glucose suggests that the glucose specifically interacts with B8CYA8 in solution.

**Figure 2.**
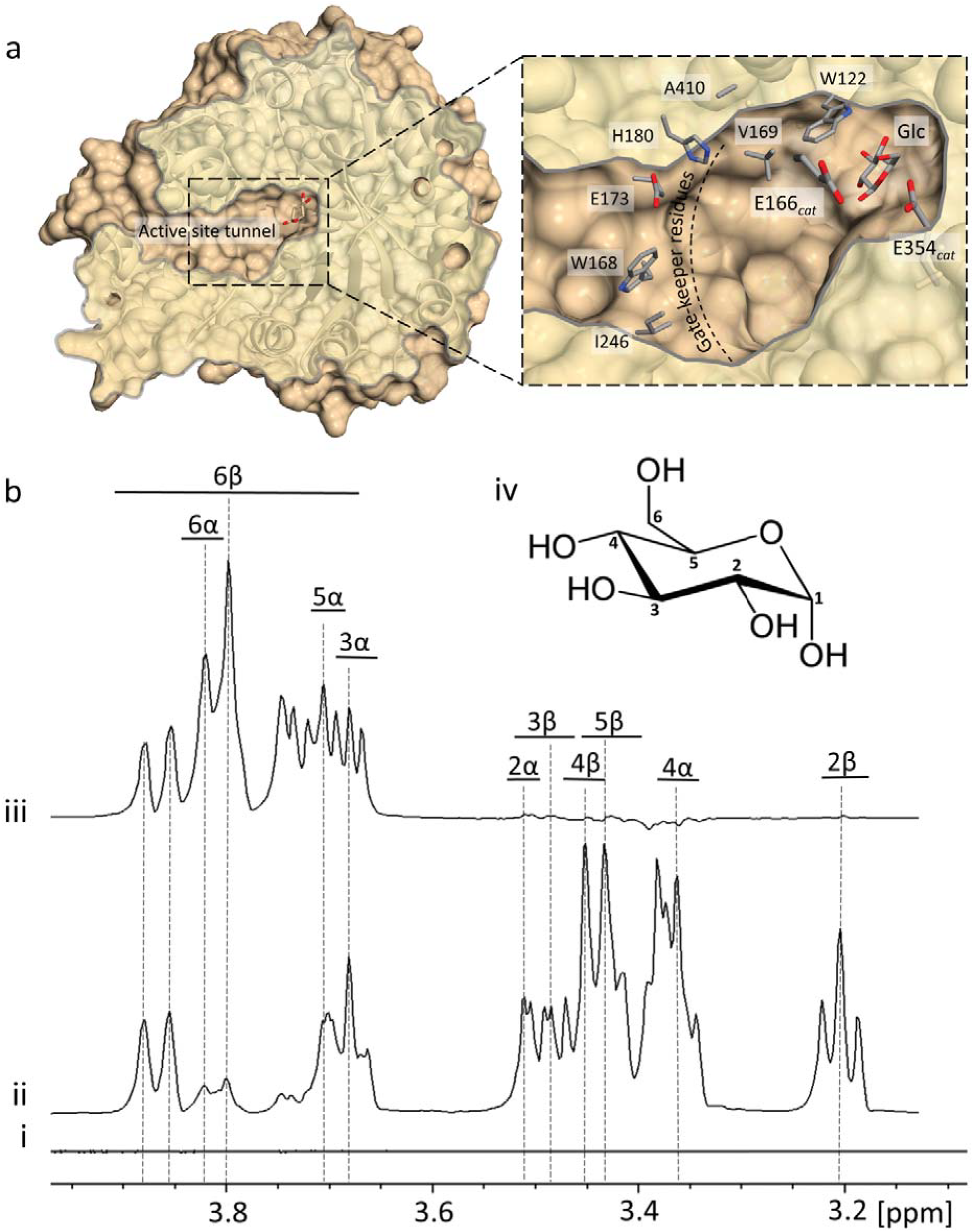
a: Cross-section of B8CYA8 (PDB 4PTX) active site pocket is shown as a space-filling model with the catalytic residues (E166 and E354), the hydrolysis product glucose, and selected gatekeeper residues (amino acid residues are shown as sticks) are labeled b: 1D-NMR spectra of glucose from (i) 1D-STD NMR experiment recorded with only glucose as control, (ii) reference 1D NMR spectrum of glucose, (iii) 1D-STD NMR spectrum of 20 mM glucose in presence of 90 μM B8CYA8. The corresponding protons in the glucose are labeled. (iv) Chemical structure of D-glucose with the labeled protons highlighted.

In the reference spectra, except for H1, all the resonances for the protons connected to the individual carbon atoms of glucose was observed. In the crystal structure of B8CYA8 complexed with glucose, the ligand-binding surface is made of predominantly aromatic amino acid residues [18]. The hydrogen atoms linked to C1, C3, C5, and C6 carbon of glucose make close contact with the residues in the ligand-binding surface (Supplementary file, Fig S1). If the glucose in solution similarly interacts with B8CYA8 as in the crystal structure, saturation of the protein would result in the efficient transfer of magnetization to the protons coupled to C1, C3, C5, and C6 carbon of glucose. Fig. 2b shows the transfer-NOE peak for the H3, H5, and H6 protons of glucose, suggesting a direct interaction between glucose and the protein. Transfer-NOE peaks for the H2 and H4 proton of glucose was not observed. While these observations suggest an agreement between glucose binding in solution and the crystal structure, the stimulation in B8CYA8 specific activity cannot be explained.

### 3.3. Glucose stimulation and inhibition is not an osmolyte effect or due to transglycosylation

It may be speculated that the glucose-induced stimulation could be due to glucose acting as an osmolyte. Therefore, we looked at the specific activity of the enzyme in the presence of another sugar, sucrose. As shown in (Supplementary file, Fig. S2), there was a slight increase in *k*_cat_ by sucrose, but there was no significant stimulation as observed with glucose. Most importantly, there was no change in *K*_*m*_ in the presence of sucrose as opposed to the 1.85 to 13-fold *K*_*m*_ increase in the presence of glucose. While an osmolyte effect cannot be ruled out in the presence of other sugars, our results indicate that glucose and sucrose do not have any significant osmolyte effects. Since an increase in enzyme activity in the presence of glucose has been previously ascribed to transglycosylation, we tried to detect the longer chain transglycosylated products in the presence of different concentrations of substrate and glucose and compared to the previously reported transglycosylation of lactose by B8CYA8 [18, 33]. As can be seen (Supplementary file, Fig. S3), no transglycosylated product was detected in the presence of glucose, and thus, we could rule out its role in the glucose tolerance of B8CYA8. To further understand the mechanism of glucose-dependent regulation, the enzyme sequence and structure was probed.

### 3.4. The basis of mutant selection

B8CYA8 is a GH1 β-glucosidase and has a typical (α/β)_8_ TIM barrel fold structure with catalytic residues (Glu166 and Glu354) located deep inside the deep active site pocket. This pocket can be binned into three regions, namely, glycone binding site (−1 subsite), aglycone binding site (+1 subsite), and gatekeeper region (Supplementary file, Fig. S4). Among β-glucosidase, the gatekeeper residues at the entrance to the deep active site pocket are mostly non-conserved (Fig. 3a). Gatekeeper residues have been suggested to play essential roles in the dynamics of the substrate influx and product efflux [13]. Thus, residues with a bulkier side chain might be expected to sterically slow down the substrate influx and efflux dynamics and affect catalysis. The hydrophilic residues at the aglycone and gatekeeper regions could help glucose stick to the active site pocket, leading to inhibition of specific activity. To test this hypothesis, we made mutations across all the regions (Supplementary file, Fig. S4) of the active site pocket, as summarized in (Supplementary file, Table S2), and shown in Fig. 3b. The size of mutated residues was compared using van der Waals volume, and the hydrophobicity was analyzed through the hydropathy index [34-36].

**Figure 3.**
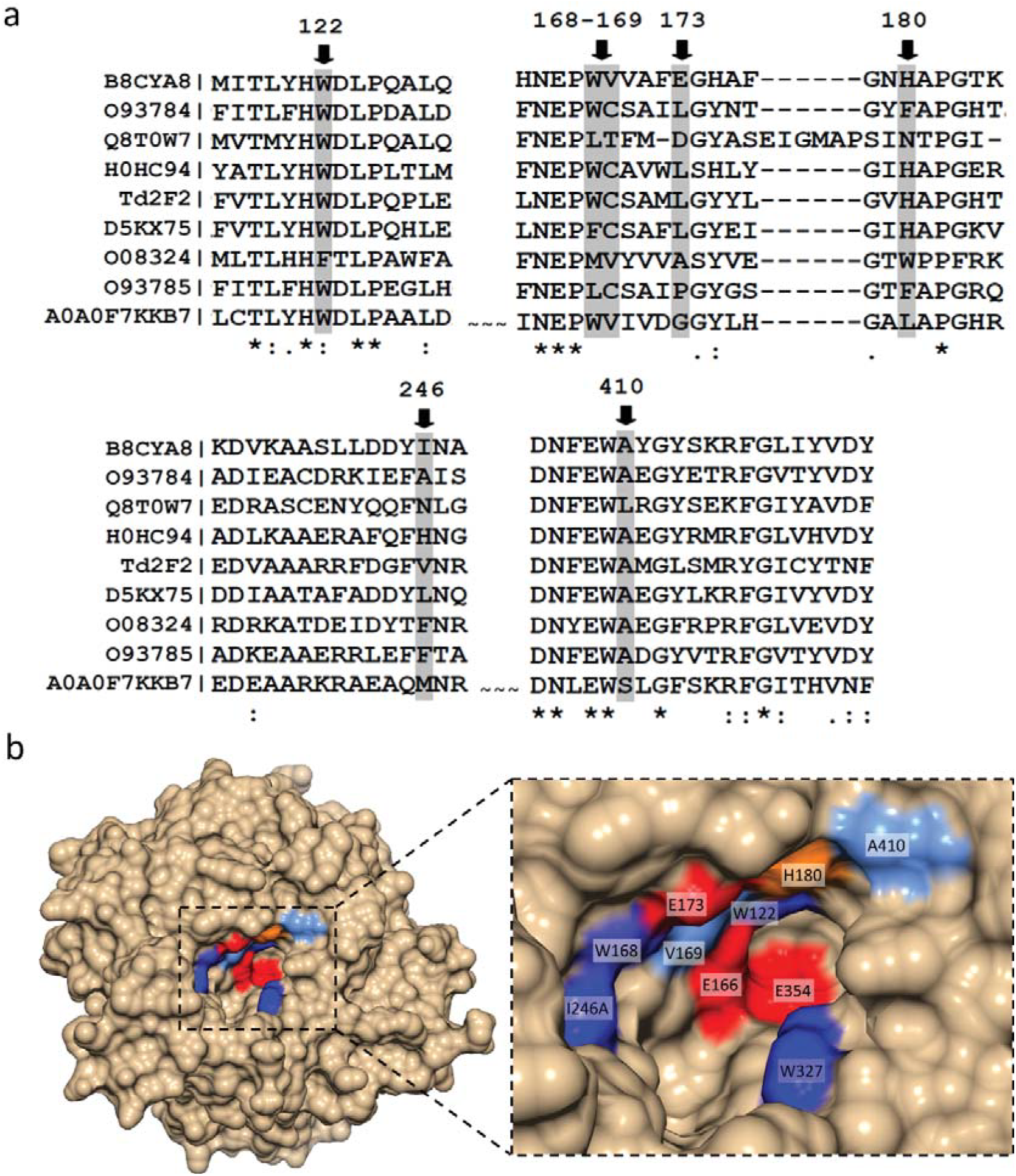
a: Multiple sequence alignment of B8CYA8 against previously identified glucose tolerant GH1 β-glucosidase by Clustal Omega [50]. B8CYA8 from *Halothermothrix orenii* [18] was aligned with O93784 from *Humicola grisea* [51], Q8T0W7 from *Neotermes koshunensis*[52], H0HC94 from *Agrobacterium tumefaciens* [22], Td2F2 from compost microbial metagenomics library [52], D5KX75 from a metagenome [53], O08324 from *Thermococcus sp.*[15] and A0A0F7KKB7 from metagenome [9]. Their UniProtKB identifies the proteins except in the case of Td2F2. b: The B8CYA8 active site pocket highlighted with the residues that were probed for a possible role in catalysis and glucose tolerance. The hydrophobic residues (W168, W327, and I246) are shown in dark blue, A410 and V169 by light blue, H180 by orange and hydrophilic residues (E166, E173, and E354) are shown in red. E166 and E354 are the catalytic acid/base and nucleophilic residues, respectively. The figure was generated using Chimera 1.10.1[54].

#### 3.4.1. Glycone binding region

The glycone binding regions in the active site pocket of β-glucosidase are typically well-conserved [37-39], as can be seen in Fig. 3a, Trp122 in the glycone binding region is conserved across most of the glucose tolerant β-glucosidase except in O08324 (the enzyme retains 100 % specific activity up to 4 M Glc) wherein a Phe is located at the equivalent position [15]. W122F mutant was constructed to understand the effect of a further increase in hydrophobicity at the glycone-binding region.

#### 3.4.2. Aglycone binding region

At the aglycone region, the residue equivalent to V169 in glucose tolerant β-glucosidase is alternately occupied by Cys and Val (Fig. 3a). We had previously reported that the substitution of Val to Cys increased B8CYA8 specific activity 1.7-fold [14]. However, glucose tolerance of wild-type or the mutant had remained unexplored.

#### 3.4.3. Gatekeeper region

The gatekeeper residues W168, E173, H180, I246, and A410 were selected to understand the effect of hydrophobicity and steric by substitution with smaller side-chain, polar side-chain and hydrophobic side-chain residues (Fig. 2a, Fig. 3b).

### 3.5. Effect of mutations on B8CYA8 specific activity in the absence of glucose

In order to determine the kinetics of the mutants on the chromogenic substrate *p*NPGlc and natural substrate cellobiose, the temperature and pH optima (T_opt_, pH_opt_) of the mutants on each of the substrates were measured (Table 1). All the mutants showed small changes in pH_opt_ in the range of 0.2 −1, and a 2 to 5 °C change in T_opt_ in comparison to the wild-type (Table 1). These subtle differences may be due to the location of mutations in the active site pocket, with small changes in interaction with the solvent molecules leading to change in pH_opt_ and similar changes in interaction with substrate molecules leading to small changes in T_opt_ [40, 41]. W122F, V169C, E173L, E173A, H180F, I246A, A410F, and A410K showed higher turnover with *p*NPGlc (Supplementary file, Table S1). While the higher specific activity of V169C, I246A, and V169C/I246A mutants was previously reported [14], the turnover numbers of A410K, V169C/E173L, and V169C/E173L/I246A increased by 17 %, 25 %, and 116 % respectively compared to wild-type B8CYA8 (Table 2).

**Table 1.**
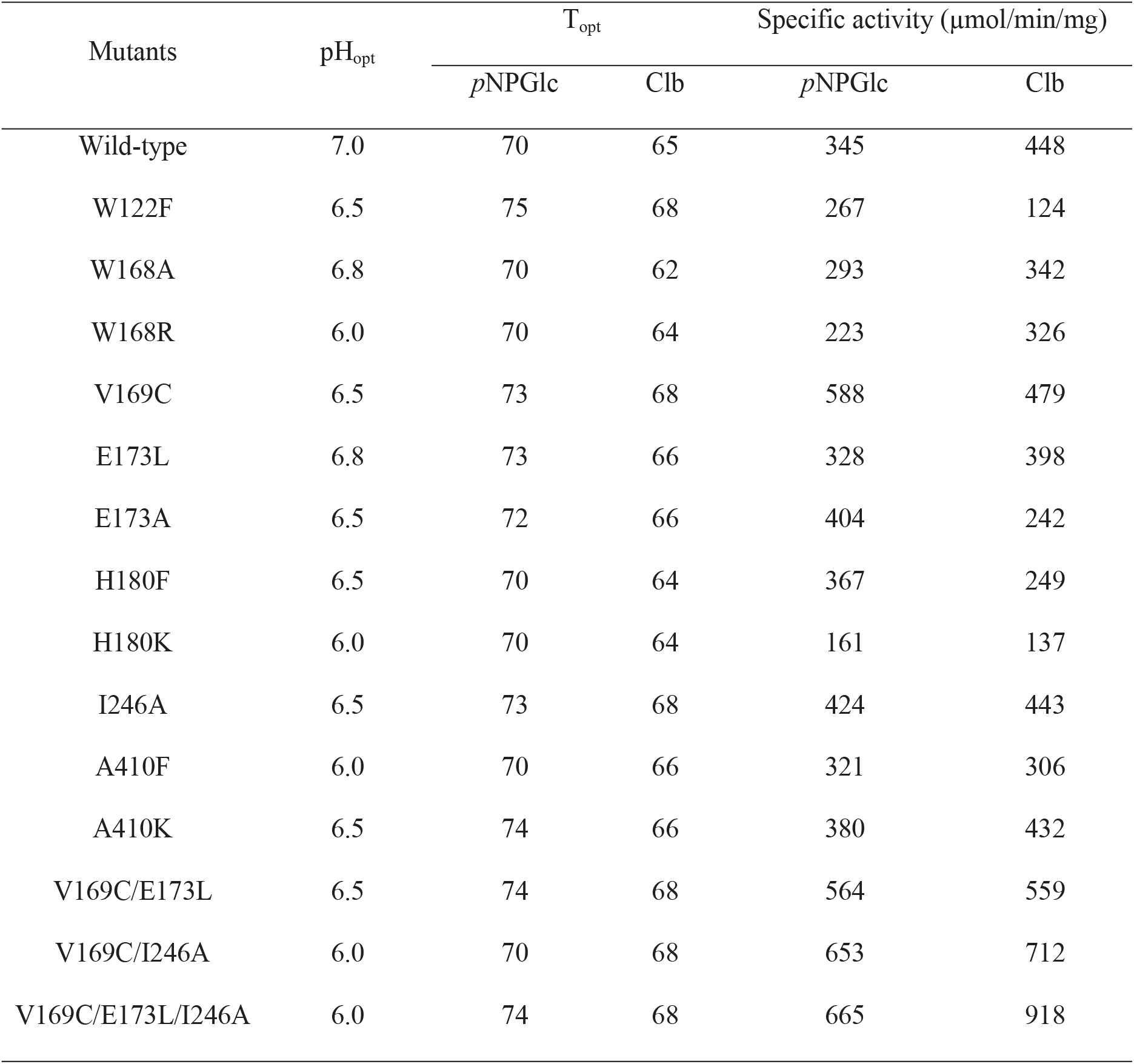
Specific activity (μmol/min/mg), temperature optimum, T_opt_ (°C) and pH optimum, pH_opt_ of recombinant wild-type B8CYA8 (Wild-type) compared to the listed mutants on *p*-nitrophenyl-D-glucopyranoside (*p*NPGlc) and cellobiose (Clb). For specific activity measurement saturation concentration of substrate was used for each mutant.

**Table 2.**
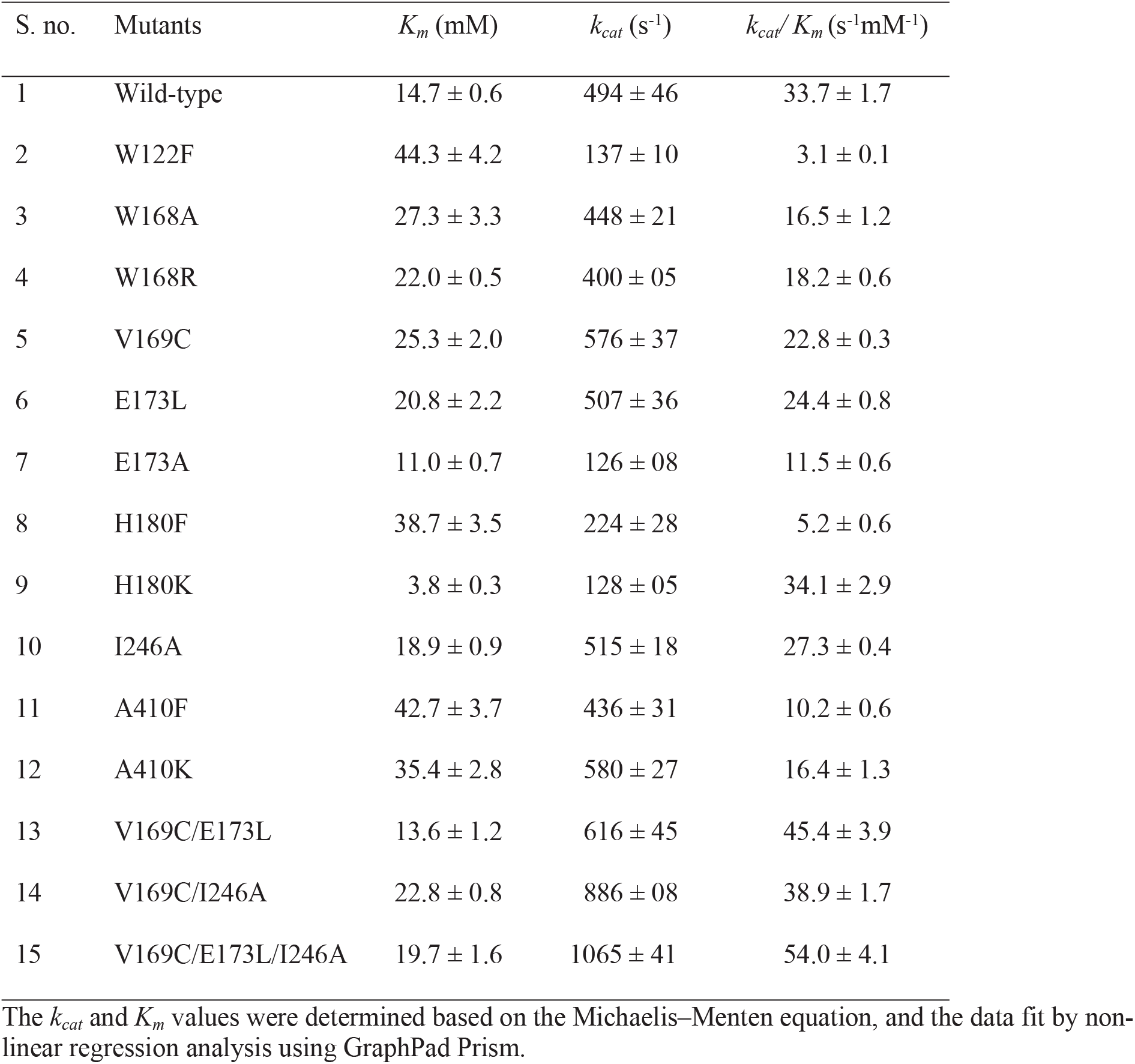
Steady-state kinetic parameters of B8CYA8 (Wild-type) and mutants on cellobiose. All measurements were in triplicates and repeated at least thrice.

### 3.6. Effect of mutations on B8CYA8 specific activity in the presence of glucose

The specific activity and kinetics of the B8CYA8 mutants were measured in the presence of exogenously added glucose (Supplementary file, Fig. S5). The direct interaction of the glucose with the enzyme possibly affects its *K*_*m*_. This variation of *K*_*m*_ of the mutants in the presence of 0 - 1.5 M glucose allowed us to bin the B8CYA8 mutants across two groups. In the first group, we considered mutants wherein we saw an increase in the fold-change in *K*_*m*_ and a decrease in glucose tolerance (Fig. 4a), and in the second group (Fig. 4b), we bin mutants wherein the glucose tolerance increased as reflected in the decrease in fold-change of *K*_*m*_ upon comparison to the wild-type. Thus, W168A/R, H180K, I246A, and A410K (Fig. 4a) showed decreased glucose tolerance while W122F, E173L/A, H180F and A410F (Fig. 4b) show increase in tolerance and stimulation (Supplementary file, Table S1). While a pattern of a decrease in fold-change of *K*_*m*_ and increase in tolerance when the residues were replaced by a more hydrophobic amino acid (Fig. 4b) is evident, the pattern of increase in fold-change in *K*_*m*_ in Fig. 4a is less so. The enzyme specificity (*k*_*cat*_/*K*_*m*_) of the improved variants are shown in Fig. 5a. Let us now look at the effect on the different regions of the active site pocket.

**Figure 4.**
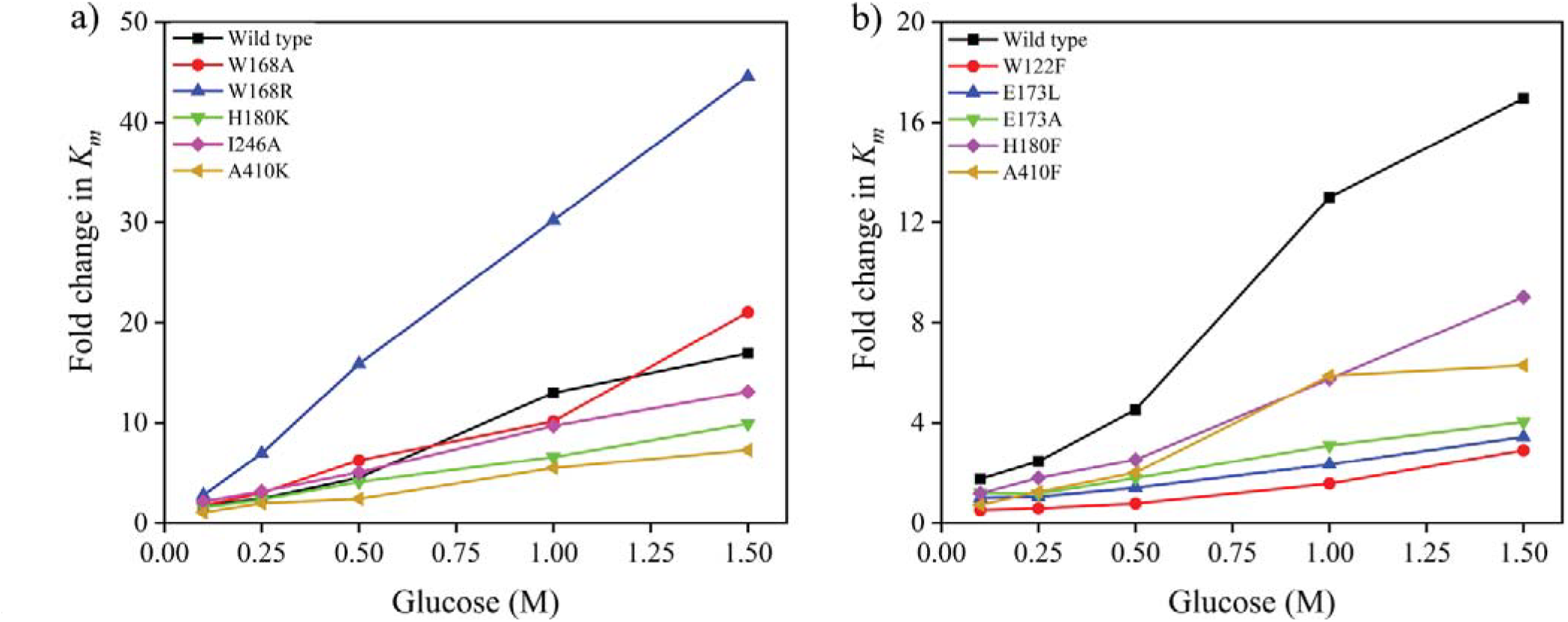
The fold change in *K*_*m*_ amongst B8CYA8 mutants vary with an increase in glucose concentration and can be binned into two groups. a) Mutants with reduced hydrophobicity and b) Mutants made more hydrophobic. The *K*_*m*_ of each mutant in the absence of glucose was normalized to 1. The enzyme kinetics were determined using the substrate, *p*NPGlc. The kinetic assay details are included in the materials and methods section.

**Figure 5.**
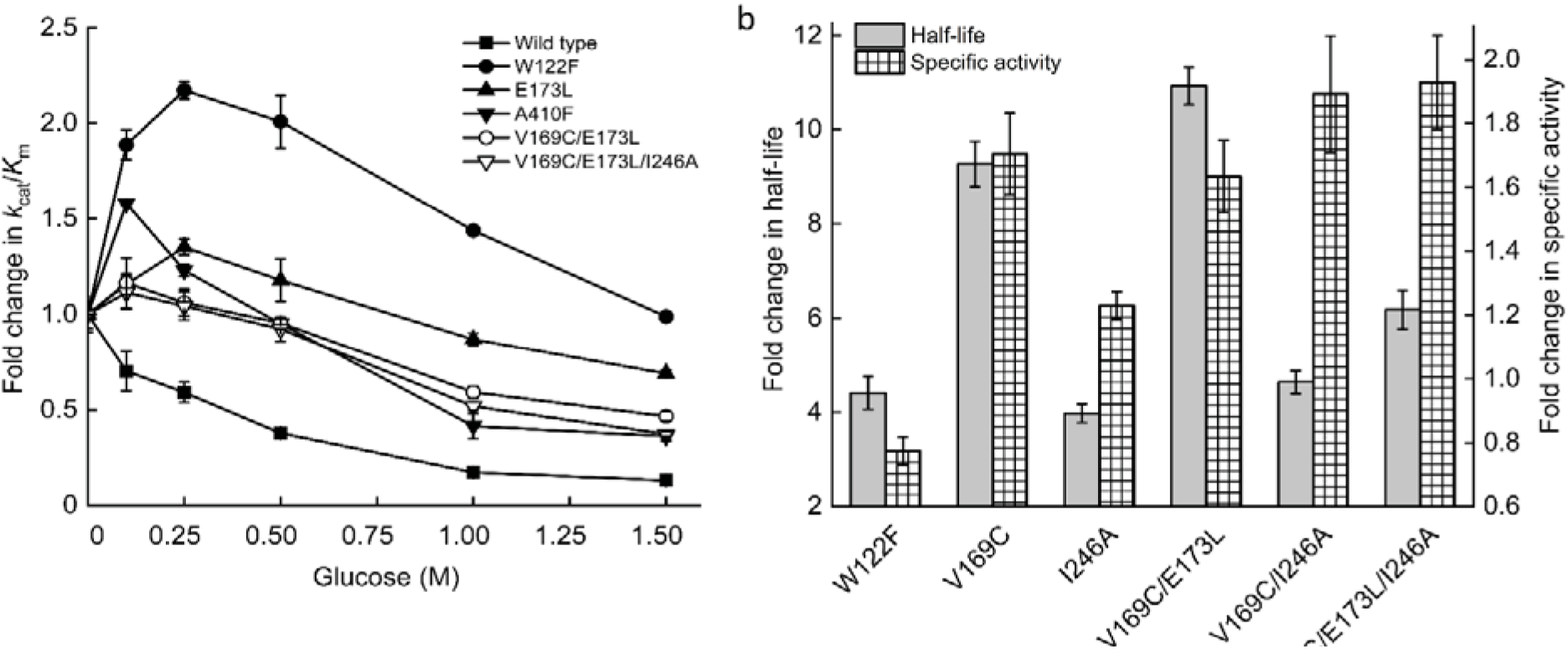
a) The fold increase in enzyme specificity (*k*_*cat*_/ *K*_*m*_) of five mutants of B8CYA8 with increasing glucose concentrations, compared to wild-type. The *k*_*cat*_/*K*_*m*_ of each mutant in the absence of glucose was normalized to 1. b) Fold change increase in half-life (Left Y-axis) of the improved mutants at 70 °C in over wild type and fold change in specific activity (Right Y-axis) under the optimum condition of each of the mutants.

#### 3.6.1. Effect of glucose at the gatekeeper region

Amongst the gatekeeper residues, W168R is drastically inhibited, as seen by a 44-fold increase in *K*_*m*_ at 1.5 M glucose. This increase in *K*_*m*_ in the presence of glucose is much higher compared to only a 17-fold increase in the wild-type. As a result, the *k*_*cat*_/*K*_*m*_ of the mutant decreased 27-fold compared to only 7.6-fold of the wild type. Mutation to a smaller and hydrophobic Ala in W168A also led to an increase in glucose inhibition, as seen from a comparatively smaller increase in *K*_*m*_ and *k*_*cat*_/*K*_*m*_. At position 173, E173L and E173A were constructed to increase hydrophobicity, and both showed an increase in glucose tolerance. The *K*_*m*_ fold change was only 3.4 and 4.0 at 1.5 M glucose, respectively, and much less than in wild-type. The reduction in enzyme efficiency, *k*_*cat*_/*K*_*m*_, was only 1.4-fold for E173L and 1.9-fold for E173A at 1.5 M glucose. In H180F, substitution by the more hydrophobic Phe increased glucose tolerance even higher, with 100 % specific activity retained at 2 M glucose. The H180K mutant showed no stimulation, and its specific activity was only 40 % at 2 M glucose. The *k*_*cat*_/*K*_*m*_ of H180F decreased only 3.6-fold while the decrease for H180K was nearly 10-fold at 1.5 M glucose. In the I246A mutant, where the hydrophobicity and residue size was decreased, less stimulation and higher inhibition were observed. Here the sterics probably play a more significant role in glucose inhibition.

#### 3.6.2. Effect of Glc at the aglycone binding site

At the aglycone binding site, the mutant V169C show negligible stimulation in the presence of glucose and its specific activity start decreasing beyond 0.75 M glucose, leading to inhibition at low glucose concentration.

#### 3.6.3. Effect of Glc at the glycone binding site

At the conserved glycone binding site, *K*_*m*_ of W122F in the absence of exogenous glucose is 5-fold higher than in WT, but in the presence of glucose, the apparent *K*_*m*_ first decreases and then starts to increase with glucose concentrations with a net 3-fold increase in *K*_*m*_ at 1.5 M glucose.

To verify that the activity and stability increases in the single mutants were additive, the V169C/E173L double mutant and V169C/E173L/I246A triple mutant was constructed. Though the initial *K*_*m*_ of the combined mutants V169C/E173L and V169C/E173L/I246A is high, there is only a 3-fold change in *K*_*m*_ in the presence of 1.5 M glucose and a 50 % increase in *k*_*cat*_/*K*_*m*_. Both mutants show higher specific activity, glucose tolerance, and kinetic stability than wild-type.

### 3.7. Effect of glucose on half-life and thermostability

The half-life of most of the mutants as well as the residual specific activity upon incubation in 1 M glucose (for 24 h at 70 °C) showed an increase in kinetic stability compared to wild-type (Supplementary file, Table S3). Notably, I246A in the presence of 1 M glucose retained more than 60 % of its specific activity after 24 h (Supplementary file, Table S3). The double and triple mutant containing V169C mutation was highly active, together the V169C/E173L/I246A had the highest increase in residual specific activity, almost 1.9-folder higher at 1 M glucose, compared to wild-type (Supplementary file, Table S3) and (Fig. 5b) These observations are in line with our previous reports when we showed that the β-glucosidase reaction product glucose improved the half-life and the kinetic stability of β-glucosidase, H0HC94 [22] and O08324 [15]. B8CYA8 and all its mutants show elevated melting temperature (Supplementary file, Table S4) with increasing concentrations of glucose, as reported previously [15, 22]. The increase was about 4-5 °C in the presence of the 0.5 M glucose, and 6-7 °C in 1 M glucose (Supplementary file, Table S4), and indicated the benefits of glucose accumulation during large-scale high biomass loading saccharification reactions.

### 3.8. Computational studies on the effect of glucose

To further understand the role of enzyme dynamics in glucose tolerance, if any, we looked at the average temperature factor (B-factor). The higher overall B-factor of glucose bound to B8CYA8 (4PTX) compared to thiocellobiose (4PTV), or 2-deoxy-2-fluoro-α-D-glucopyranose (4PTW) bound enzyme suggests that addition of glucose introduces enhanced structural flexibility to B8CYA8 (Fig. 6a). Molecular Dynamics (MD) simulations (Fig. 6 b,c,d) confirm the increased glucose-dependent backbone dynamics of the active site residues and flexibility of the active site pocket to accommodate glucose. The gatekeeper residues selected for calculation was based on the symmetry of active site pocket entrance and hence were not all similar to the sites selected for mutagenesis. Fig. 6c highlights the increase in RMSF across residues in the gatekeeper and aglycone bindings site while Fig. 6d shows the increase in backbone flexibility of residues in the glycone binding site as well as a few residues in the gatekeeper region of the active site pocket. Such flexibility could enable a glucose-induced modulation of dynamic equilibrium in the active site pocket width. Indeed, when we compared the solvent-accessible surface area (SASA) from molecular dynamics simulation trajectories of the gatekeeper residues (residues 299, 314, 316, 324, 325, 326, 410 and 411) between 0.05 M and 1.5 M glucose, we observed an increase in the distribution of total surface area with increasing glucose concentration (shown in Fig. 6e). The SASA for the eight individual residues mentioned above is shown in Fig. 6f.

**Figure 6.**
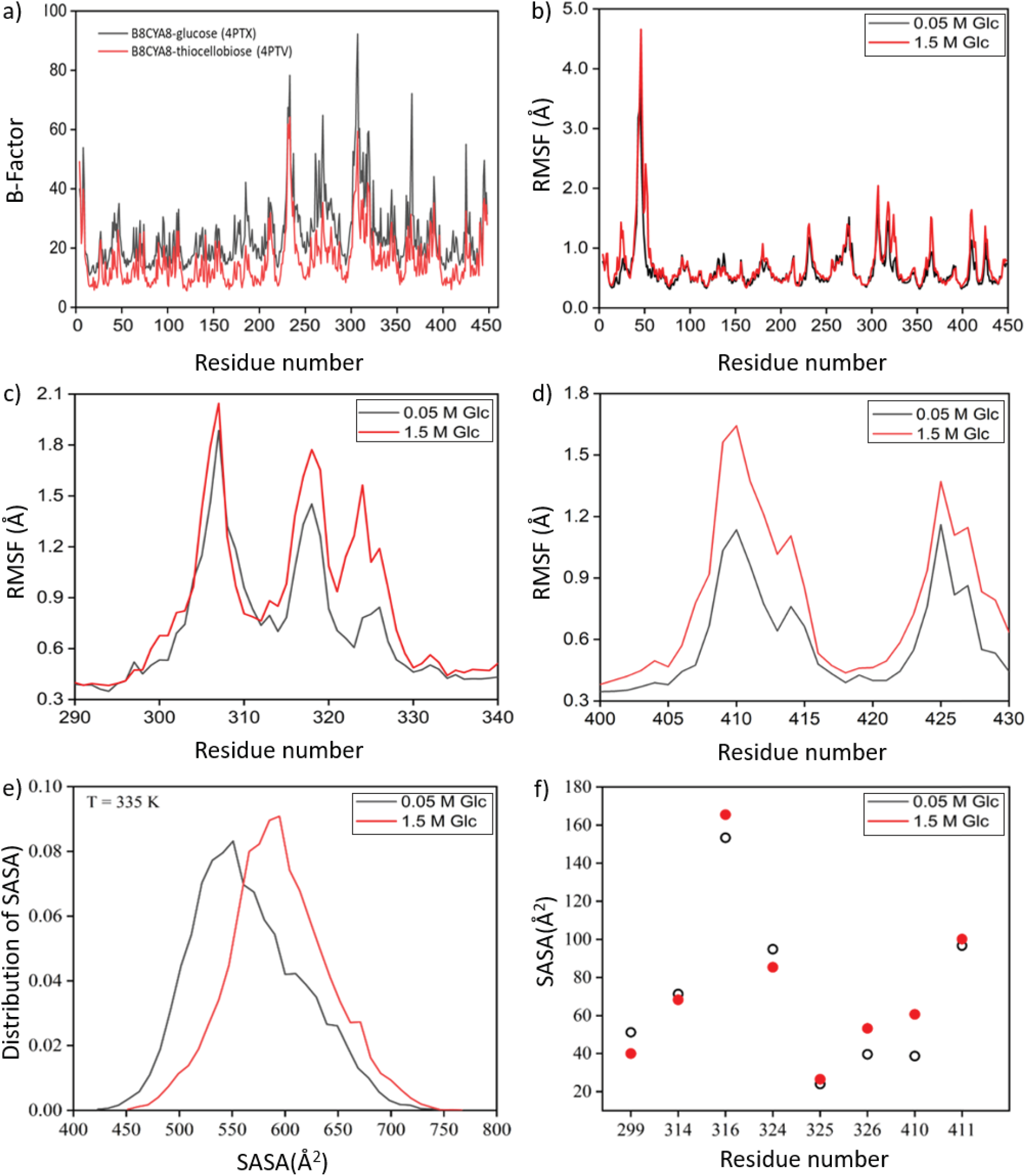
The effect of glucose on protein dynamics. a) The B-factor comparison of the B8CYA8 crystal structure (4PTX: B8CYA8 with glucose; 4PTV: B8CYA8 with thiocellobiose) shows the fluctuation differences of residues. B8CYA8-glucose complex structure (4PTX) has higher residue fluctuations than B8CYA8-thiocellobiose (4PTV) complex. b) RMSF of all B8CYA8 residues (top) at T = 335 K in the presence of 0.05 M (black) and 1.5 M (red) glucose (Glc). (c) The RMSF of residues 290-340 and (d) residues 400-430 highlight the increase in RMSF of gatekeeper, glycone, and aglycone binding residues in the active site pocket. (e) Average SASA (solvent accessible surface area) distribution of the gate-keeper residues (residues 299, 314, 316, 324, 325, 326, 410 and 411) at 335 K temperature in the presence of 0.05 M and 1.5 M glucose. (f) SASA of the individual gate-keeper residues. The structures in (a) were taken from the Protein Database [18]. The structures for analyses (b-f) were taken from the MD simulation trajectories.

### 3.9. Synergy with commercial cellulase

The synergistic effects of B8CYA8 and mutants (V169C, V169C/E173L, V169C/I246A, and V169C/E173L/I246A) on the model substrate Avicel was evaluated by assaying in the presence of the commercially available *T. viride* cellulase cocktail (Supplementary file, Table S5). A commercially available sweet almond β-glucosidase was used as a control. Saccharification supplementation by the triple mutant, V169C/E173L/I246A, showed a 90 % increase in glucose yield compared to only *T. viride* cellulase. The chosen reaction condition (pH 5 and 37 °C) was optimum for only the sweet almond β-glucosidase and *T. viride* cellulase and in spite, the B8CYA8 and its mutants contributed to the increased saccharification efficiencies. A further improvement in glucose yield upon B8CYA8 triple mutant supplementation can be expected upon use as part of a thermophilic cocktail optimized for activity at similar T_opt_ and pH_opt_.

## 4. Discussion

Previously we reported the role of mutations in the non-conserved residues, in the active the pocket of B8CYA8, V169C, and E173L, towards engineering higher catalytic efficiencies and thermal stability [14]. Here we investigated the effect of glucose on the catalytic efficiency of the enzyme and the role of the active site pocket. We observed that the specific activity of B8CYA8 on the chromogenic substrate *p*NPGlc increases with glucose concentration, resulting in an increase in *k*_*cat*_ and the apparent *K*_*m*_. This increase in activity in the presence of glucose has been previously ascribed to transglycosylation [33]. Since we did not observe any transglycosylated products, we ruled out transglycosylation as a factor in the glucose tolerance of B8CYA8. Our studies with sucrose rule out osmolyte effects as a significant factor in glucose stimulation. Glucose has been conjectured to inhibit β-glucosidase by direct binding to the active site and compete with the substrate. The STD NMR study provided substantial evidence of the direct interaction of H3, H5 and H6 hydrogen of glucose with B8CYA8 residues, and together with the crystal structure suggest that glucose can specifically interact with the protein in the solution and bind to a region/subsite inside the active site pocket such that despite glucose binding to the active site, the enzyme is initially stimulated. It may, however, be possible that there are other low-affinity binding sites on the protein.

Kinetic data of B8CYA8 and its mutants in the presence of exogenously added glucose reveal the importance of the gatekeeper residues in glucose entry and interaction in the active site pocket. The large non-polar side chain of Trp, Leu, Phe, and Ile (W168, E173L, H180F, and I246) along with the strong hydrophobic interaction affects the entry of glucose inside the active site pocket. Conversely, polar side chains at the gatekeeper region (W168R, E173, and H180K) facilitate the necessary electrostatic interactions for the accommodation of glucose near the entrance of the pocket and competitively inhibits the enzyme. Residues with smaller side-chain (W168A and I246A) enable glucose entry inside the pocket, and in turn the inhibit enzyme activity. At the aglycone binding (+1 subsite) site, V169C specific activity is the highest among the mutants (1.8-fold increase in k_cat_ in the absence of glucose, compared to the WT) such that the addition of exogenous glucose (1 M glucose) does not increase the specific activity. The polar side chain probably changes the geometry of the hydrogen bonding network at the catalytic sites to facilitate glucose accumulation. At the glycone binding site, W122F showed increased glucose tolerance, as in the previously reported β-glucosidase (O08324) in *Thermococcus sp.*[15]. Here the indole ring of Trp probably provides a geometrically complementary apolar surface for interaction with glucose, and its π-electron cloud favorably interacts with the positively charged aliphatic protons of glucose [42]. The Phe may prevent the accumulation of glucose near the active site from enhancing the glucose tolerance of W122F. Similar mutations across H0HC94 and O08324 studied in our laboratory, seems to support the role of hydrophobic residues inside the active site pocket [12, 22]. Amino acid residues in the aglycone-binding site have been proposed to be responsible for glucose tolerance [13, 43]. Based on *in-silico* docking studies, Yang et al. proposed that glucose tolerant β-glucosidase have a higher propensity of glucose binding near the middle of the pocket while less tolerant one can have glucose binding at the bottom [44]. The B8CYA8 crystal structure in the presence of glucose and STD-NMR studies, however, show the interaction with glucose near the bottom of the active site pocket. Glucose inhibition has been attributed to binding to other allosteric sites, and by non-productive binding to other sites in the active site pocket [11, 45] However cooperativity in B8CYA8 could not be established by measurement of hill coefficient (Supplementary file, Table S6) which were around one in the wild-type and the mutants. The binding of glucose to secondary binding site(s) is, however, yet to be experimentally proven.

The higher average temperature factor (B-factor) of glucose bound to B8CYA8 (4PTX) compared to the substrate or inhibitor-bound structures suggested that the addition of glucose introduces enhanced structural flexibility into B8CYA8. Such a trend was also confirmed upon observation of higher B-factors of two other glucose tolerant β-glucosidase structures in the presence of glucose, in comparison with the respective structures in the absence of glucose [33, 46, 47]. MD simulations confirmed that glucose increased the backbone dynamics of the gatekeeper, glycone and aglycone binding site residues and flexibility of the active site pocket to accommodate glucose. A flexible active site would enable a glucose-induced modulation of dynamic equilibrium in the active site pocket width. Indeed, the solvent-accessible surface area (SASA) of selected gatekeeper residues measured from molecular dynamics simulation trajectories at 0.05 M and 1.5 M glucose show an increase in the distribution of total surface area with increasing glucose concentrations. Thus, glucose may stabilize a widened active site pocket structure and facilitate substrate accessibility to the active site.

Previously it was reported that the addition of small amounts of glucose could reduce the non-productive binding of substrate to +1 and +2 subsites and stimulate enzyme activity [45]. A comparison with PDB structure 3F5K and 2O9P indicates the presence of the +2 subsite in B8CYA8 which can potentially assist in the non-productive binding of the substrate. Recently we showed that in the presence of low substrate concentrations (1 mM *p*NPGlc), B8CYA8 is inhibited at all concentrations of glucose [48]. We also reported by MD simulations of B8CYA8 in the presence of glucose that at the gatekeeper region, the number of glucose molecules increases significantly with glucose concentration than inside the pocket [48]. Our results reported here do not preclude the possibility of substrate binding non-productively to the +1 to +2 subsite or the possibility of glucose binding at this subsite to relieve the non-productive binding of the substrate and increase enzyme activity. At higher substrate concentrations, low concentrations of glucose may relieve the enzyme from the non-productive binding to increase enzyme activity, in addition to the active site pocket widening. Thus, the process of glucose-induced pocket broadening and stimulation and inhibition seems to be highly dynamic and sensitive towards the ratio of the substrate and glucose. Fig. 7 shows our proposed model summarizing the role of hydrophobicity, size, and glucose on the active site pocket. In the absence of glucose, the substrate molecule dominates the inside of the pocket due to interactions with the hydrophilic residues and may bind to the non-productive binding sites inside the pocket. In the presence of high concentrations of glucose, the active site pocket broadens and may facilitate the accumulation of substrate and glucose inside the pocket. Active site pockets lined with hydrophobic residues would be expected to discourage glucose binding and therefore decrease the number of glucose molecules inside the active site pocket. The molecular basis of the glucose-induced active site pocket dynamics at different substrate concentrations is currently under investigation by simulating the enzyme behavior in the presence of different concentrations of substrate and glucose to understand this complex interplay.

**Figure 7.**
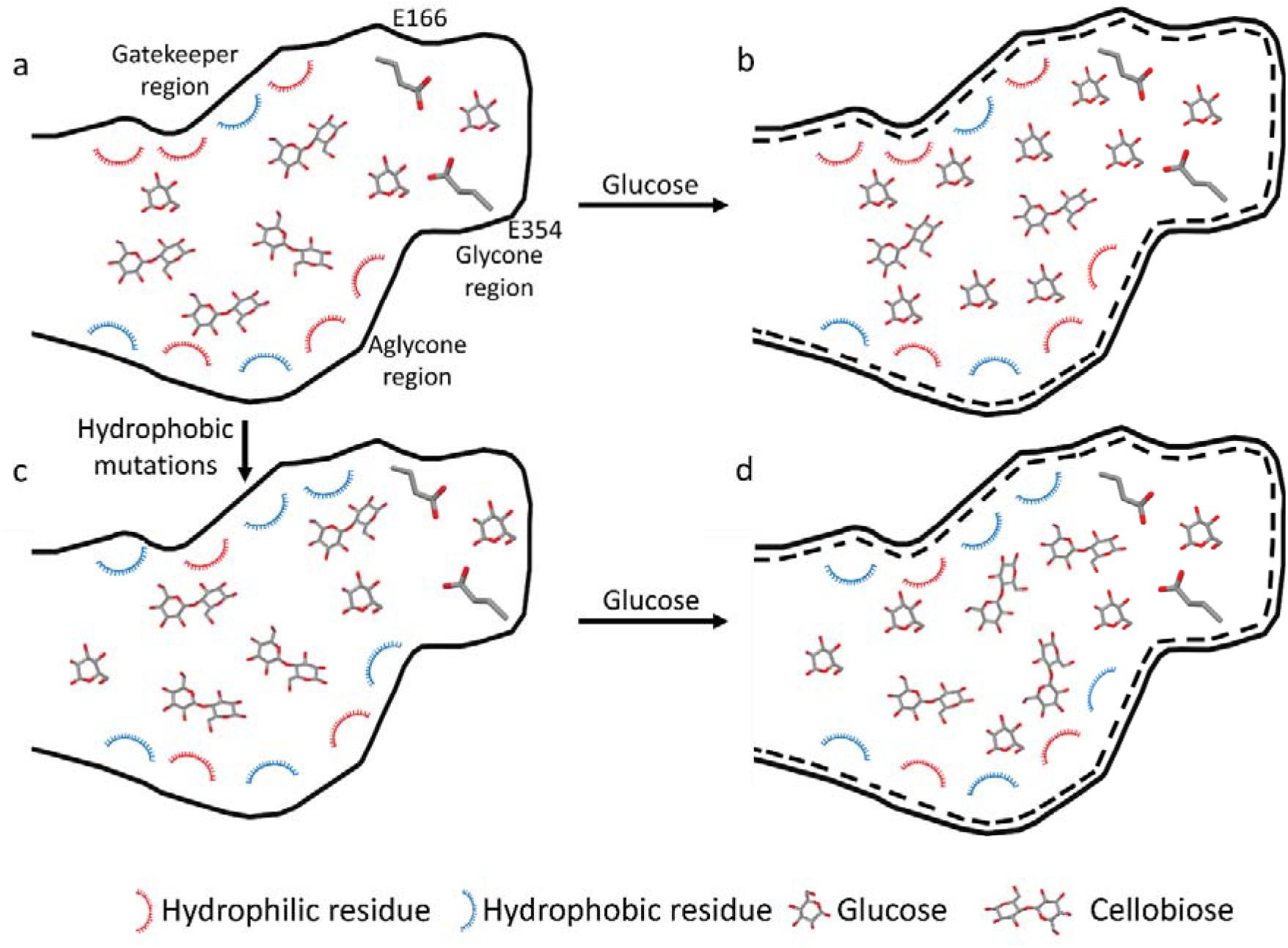
Schematic of the effect of glucose on active site pocket of B8CYA8. a) Active site pocket with hydrophilic residues in the gatekeeper region of the enzyme. In the absence of glucose, the substrate molecule dominates the inside of the pocket due to interactions with the hydrophilic residues. b) Excess of glucose broadens the active site pocket and increases the accumulation of glucose inside the pocket, along with more substrate and leads to competitive inhibition of the substrate. c) When the active site pocket has more hydrophobic residues (by mutations at gatekeeper region or in the wild-type for glucose tolerant β-glucosidase). d) Upon addition of glucose to the enzyme with hydrophobic residues at the gatekeeper region, an excess of glucose increases the pocket width, but glucose cannot stick around inside the pocket due to the greater number of hydrophobic residues and leading to an increase in glucose tolerance/ decrease in glucose inhibition.

Another essential objective of this study was to engineer improved variants of β-glucosidase towards a thermophilic cellulase cocktail. The triple mutant V169C/E173L/I246A is particularly valuable, with a three-fold increase in turnover number on natural substrate cellobiose (*k*_*cat*_ = 1065 s^−1^), long half-life of more than 7 hours at 70 °C, high residual specific activity of around 75 % after a 24 h incubation in 1.0 M glucose at 70 °C. Our initial studies on the model substrate Avicel support the potential gains of using a high specific activity and glucose tolerant β-glucosidase [14]. We had also previously reported the potential of recycling the wild-type enzyme towards industrial applications [49].

In summary, we report that B8CYA8 exhibits both stimulation and inhibition by glucose that is not due to transglycosylation or osmolyte effects. The increase in hydrophobicity inside the active site pocket probably increases substrate as well as product accessibility, and the presence of high glucose concentrations modulate the tunnel width. While our studies do not rule out the possibility of non-productive substrate-binding playing a role, the dynamic modulation of the active site pocket by glucose and substrate seems to dictate stimulation or inhibition of enzymatic activity. Our studies reveal the role of non-conserved residues in the active site pocket and the benefits of engineering such residues.

## 5. Acknowledgments

This work was supported in part by the Science & Engineering Research Board (SERB), Government of India, EMR/2016/003705 (S.D.), Energy Bioscience Overseas Fellowship, Department of Biotechnology, Government of India, BT/NBDB/22/06/2011 (S.D), and Academic Research Fund (IISER Kolkata) (SD, RD and PKG). A Senior Research Fellowship from CSIR supports SKS, and SDas was supported by an Inspire Fellowship, DST, Govt. India. SK is supported by an Institute Fellowship from IISER Kolkata. We thank Mr. Shubhasish Goswami for his initial help in cloning.

## 6. Author contributions

SD, SKS, SDas, RD, and PKG designed the study. SKS, SDas, SK, and RD performed the study. SD, SKS, SDas, RD, PKG analyzed the data. SD, SKS, and RD wrote the paper.

## 7. Compliance with Ethical Standards

Conflict of Interest: All authors declare that they have no conflict of interest.

Ethical Approval: This article does not contain any studies with human participants or animals performed by any of the authors.

