## Supplemental File for "Elucidating the regulation of glucose tolerance through the interaction between the reaction product and active site pocket residues of a β-glucosidase from *Halothermothrix orenii*"

[1]Protein Engineering Laboratory, Department of Biological Sciences, Indian Institute of Science Education and Research Kolkata, Mohanpur, India and [2]Department of Chemical Sciences, Indian Institute of Science Education and Research Kolkata, Mohanpur, India and [3]Department of Biological Sciences, Indian Institute of Science Education and Research Kolkata, Mohanpur, India and [4]Centre for Advanced Functional Materials, Indian Institute of Science Education and Research Kolkata, Mohanpur, India and [5]Centre for Climate and Environmental Sciences, Indian Institute of Science Education and Research Kolkata, Mohanpur, India

Methods:

***Differential scanning fluorimetry (DSF) to determine melting temperature, $T_m$:*** DSF Analysis was performed as per protocol previously described, wherein the unfolding of protein was recorded by fluorescence intensity versus temperature plot and the data transformed and analyzed [1, 2]. Each measurement was made in triplicate.

***Thin Layer Chromatography (TLC):*** TLC was performed to analyze the hydrolysis product catalyzed by B8CYA8. Preactivated Kieselgel 60 F-254 silica gel plate (Merck Millipore, Mumbai, India) along with eluent: n-butanol–n-propanol–ethanol-water (2:3:3:2) was used for the detection of products. The samples were taken from activity assays measurement at 70 °C in the presence of different concentrations (0-1.5 M) of glucose and 0.1 M lactose as control.

Table S1. Kinetic parameters of B8CYA8 mutants in the presence of different concentrations of glucose with substrate *p*NPGlc. Kinetic parameters were determined in optimized reaction conditions for each mutant, as described in Table 1. Final glucose concentrations of 0.1 M, 0.25 M, 0.5 M, 1.0 M, and 1.5 M in a final reaction mixture of 100 µL were used. Experiments were done in triplicate and repeated at least thrice. Errors shown here are standard deviations of independent reactions. Already published data [3] are shown in *italics*.

| **Mutants** | **Kinetic parameter** | **Glc (M)** | | | | | |
|---|---|---|---|---|---|---|---|
| | | **0** | **0.1** | **0.25** | **0.5** | **1** | **1.5** |
| Wild type | $K_m$ (mM) | *1.8 ± 0.1* | 3.2 ± 0.3 | 4.4 ± 0.6 | 8.1 ± 0.3 | 23.4 ± 2.2 | 30.5 ± 3.8 |
| | $k_{cat}$ ($s^{-1}$) | *339.0 ± 28.3* | 411.5 ±16.1 | 482.2 ± 28 | 572.0 ± 5.3 | 752.7 ± 22.8 | 742.8 ± 49.5 |
| | $k_{cat}/K_m$ ($s^{-1}mM^{-1}$) | *185.5 ± 3.3* | 130.4 ±19.2 | 109.7 ± 9.8 | 70.2 ± 1.9 | 32.3 ± 4.0 | 24.5 ± 1.5 |
| W122F | $K_m$ (mM) | 9.5 ± 0.1 | 4.9 ± 0.2 | 5.6 ± 0.4 | 7.5 ± 0.4 | 15.0 ± 0.4 | 27.6 ± 1.4 |
| | $k_{cat}$ ($s^{-1}$) | 359.8 ± 23.6 | 390.6 ± 33.2 | 499.2 ± 43.9 | 626.3 ± 12.0 | 904.1 ± 26.2 | 1144.7 ± 50.4 |
| | $k_{cat}/K_m$ ($s^{-1}mM^{-1}$) | 41.9 ± 3.9 | 79.0 ± 3.3 | 90.9 ± 1.9 | 84.1 ± 5.8 | 60.2 ± 0.0 | 41.4 ± 0.6 |
| W168A | $K_m$ (mM) | 1.9 ± 0.2 | 3.4 ± 0.1 | 5.7 ± 0.4 | 11.9 ± 1.0 | 19.3 ± 2.3 | 40.0 ± 0.04 |
| | $k_{cat}$ ($s^{-1}$) | 283.1 ± 28.2 | 345.4 ± 18.8 | 447.8 ± 5.2 | 567.3 ± 35.3 | 596.0 ± 37.3 | 678.7 ± 16.9 |
| | $k_{cat}/K_m$ ($s^{-1}mM^{-1}$) | 147..3 ± 6.4 | 102.9 ± 8.8 | 72.0 ± 7.1 | 47.7 ± 1.2 | 31.0 ± 1.9 | 17.0 ± 0.44 |
| W168R | $K_m$ (mM) | 1.4 ± 0.1 | 3.9 ± 0.3 | 9.7 ± 0.5 | 22.2 ± 2.5 | 42.3 ± 3.6 | 62.4 ± 2.4 |
| | $k_{cat}$ ($s^{-1}$) | 209.1 ± 7.82 | 272.7 ± 12.0 | 303.8 ± 20.2 | 363.1 ± 27.1 | 463.9 ± 37.0 | 334.3 ± 3.8 |
| | $k_{cat}/K_m$ ($s^{-1}mM^{-1}$) | 147.5 ± 17.7 | 70.5 ± 8.5 | 31.3 ± 4.0 | 16.5 ± 2.8 | 11.0 ± 0.3 | 5.4 ± 0.2 |
| V169C | $K_m$ (mM) | *3.5 ± 0.1* | 6.3 ± 0.3 | 8.6 ± 0.5 | 14.0 ± 1.0 | 26.2 ± 1.3 | 38.9 ± 4.6 |
| | $k_{cat}$ ($s^{-1}$) | *604.7 ± 30.3* | 828.6 ± 9.7 | 934.4 ± 34.9 | 1095.6 ± 80.0 | 1089.7 ± 108.3 | 1101.8 ± 70.3 |
| | $k_{cat}/K_m$ ($s^{-1}mM^{-1}$) | *174 ± 1.6* | 132.6 ± 3.8 | 108.8 ± 10.2 | 78.4 ± 0.1 | 41.6 ± 2.4 | 28.4 ± 1.7 |

**Continued…**

| Mutants | Kinetic parameter | Glc (M) | | | | | |
|---|---|---|---|---|---|---|---|
| | | 0 | 0.1 | 0.25 | 0.5 | 1 | 1.5 |
| E173L | $K_m$ (mM) | 3.6 ± 0.1 | 3.6 ± 0.2 | 3.5 ± 0.3 | 5.1 ± 0.6 | 8.4 ± 0.7 | 12.4 ± 0.4 |
| | $k_{cat}$ ($s^{-1}$) | 349.1 ± 12.8 | 408.3 ± 6.8 | 498.8 ± 30.9 | 582.3 ± 10.6 | 714.4 ± 34.6 | 834.9 ± 48.0 |
| | $k_{cat}/K_m$ ($s^{-1}mM^{-1}$) | 97.6 ± 0.2 | 113.0 ± 5.3 | 131.9 ± 4.0 | 115.1 ± 10.7 | 84.8 ± 3.1 | 67.4 ± 1.7 |
| E173A | $K_m$ (mM) | 7.1 ± 0.7 | 8.6 ± 0.3 | 8.4 ± 1.0 | 12.8 ± 1.3 | 22.0 ± 0.4 | 28.7 ± 1.4 |
| | $k_{cat}$ ($s^{-1}$) | 491.3 ± 18.4 | 617.2 ± 30.0 | 678.9 ± 11.8 | 832.2 ± 79 | 970.7 ± 26.8 | 1021.7 ± 12.9 |
| | $k_{cat}/K_m$ ($s^{-1}mM^{-1}$) | 69.1 ± 4.2 | 71.5 ± 1.0 | 81.7 ± 8.5 | 65.1 ± 0.2 | 44.1 ± 0.4 | 35.7 ± 1.3 |
| H180F | $K_m$ (mM) | 2.9 ± 0.0 | 3.4 ± 0.1 | 5.2 ± 0.6 | 7.3 ± 3.5 | 16.7 ± 4.2 | 26.2 ± 3.5 |
| | $k_{cat}$ ($s^{-1}$) | 388.1 ± 34.9 | 515.1 ± 43.3 | 669.6 ± 69.8 | 811.9 ± 22.3 | 848.6 ± 80.4 | 979.1 ± 89.9 |
| | $k_{cat}/K_m$ ($s^{-1}mM^{-1}$) | 132.4 ± 13.7 | 149.4 ± 8.7 | 127.7 ± 0.2 | 83.1 ± 4.1 | 57.3 ± 0.3 | 36.3 ± 2.1 |
| H180K | $K_m$ (mM) | 2.2 ± 0.2 | 3.5 ± 0.3 | 5.2 ± 0.6 | 9.0 ± 1.0 | 14.5 ± 0.3 | 21.9 ± 2.9 |
| | $k_{cat}$ ($s^{-1}$) | 154.8 ± 8.3 | 181.2 ± 3.0 | 201.6 ± 3.8 | 227.2 ± 1.3 | 223.7 ± 10.1 | 160.0 ± 13.3 |
| | $k_{cat}/K_m$ ($s^{-1}mM^{-1}$) | 71.6 ± 1.3 | 52.8 ± 6.1 | 38.9 ± 3.5 | 25.3 ± 2.5 | 15.4 ± 0.4 | 7.3 ± 0.4 |
| I246A | $K_m$ (mM) | *1.9 ± 0.2* | 4.1 ± 0.4 | 6.0 ± 0.7 | 9.6 ± 1.0 | 18.4 ± 1.3 | 24.9 ± 1.4 |
| | $k_{cat}$ ($s^{-1}$) | *415.5 ± 32.7* | 558.4 ± 37.4 | 717.5 ± 57.0 | 950.7 ± 54.5 | 915.8 ± 58.3 | 631.3 ± 18.4 |
| | $k_{cat}/K_m$ ($s^{-1}mM^{-1}$) | *218.4 ± 15.7* | 139.1 ± 4.2 | 99.4 ± 5.6 | 77.8 ± 5.6 | 41.2 ± 1.9 | 25.4 ± 1.7 |
| A410F | $K_m$ (mM) | 3.7 ± 0.3 | 2.7 ± 0.1 | 4.6 ± 0.1 | 7.5 ± 0.1 | 21.8 ± 3.1 | 23.3 ± 3.5 |
| | $k_{cat}$ ($s^{-1}$) | 343.5 ± 15.1 | 395.9 ± 15.2 | 529.0 ± 6.9 | 660.8 ± 11.2 | 831.7 ± 16 | 780.0 ± 66.5 |
| | $k_{cat}/K_m$ ($s^{-1}mM^{-1}$) | 93.0 ± 2.6 | 147.0 ± 1.5 | 114.4 ± 3.0 | 88.4 ± 2.6 | 38.7 ± 6.2 | 33.7 ± 2.1 |

**Continued…**

| Mutants | Kinetic parameter | Glc (M) | | | | | |
|---|---|---|---|---|---|---|---|
| | | **0** | **0.1** | **0.25** | **0.5** | **1** | **1.5** |
| A410K | $K_m$ (mM) | 3.0 ± 0.2 | 3.2 ± 0.2 | 6.0 ± 0.4 | 7.3 ± 0.5 | 16.5 ± 0.9 | 21.9 ± 1.3 |
| | $k_{cat}$ ($s^{-1}$) | 394.4 ± 5.0 | 509.2 ± 8.0 | 698.6 ± 41.5 | 788.7 ± 49.5 | 1019.6 ± 18.4 | 987.2 ± 22.4 |
| | $k_{cat}/K_m$ ($s^{-1}mM^{-1}$) | 133.1 ± 9.2 | 157.3 ± 6.4 | 110.5 ± 5.7 | 108.2 ± 6.8 | 61.9 ± 4.2 | 45.4 ± 1.2 |
| V169C/E173L | $K_m$ (mM) | 6.6 ± 0.5 | 6.5 ± 0.7 | 8.5 ± 1.0 | 10.5 ± 0.1 | 19.0 ± 1.7 | 22.3 ± 1.8 |
| | $k_{cat}$ ($s^{-1}$) | 670.4 ± 1.2 | 767.3 ± 1.5 | 916.1 ± 51.1 | 1027.7 ± 21.6 | 1152.7 ± 92.3 | 1062.4 ± 27.4 |
| | $k_{cat}/K_m$ ($s^{-1}mM^{-1}$) | 102.5 ± 7.6 | 119.4 ± 13.4 | 108.8 ± 7.2 | 97.9 ± 3.1 | 60.8 ± 3.3 | 47.8 ± 2.6 |
| V169C/I246A | $K_m$ (mM) | *4.19 ± 0.44* | 4.9 ± 0.2 | 8.7 ± 0.4 | 15.4 ± 1.2 | 34.6 ± 0.5 | 35.5 ± 2.4 |
| | $k_{cat}$ ($s^{-1}$) | *692.4 ± 43.67* | 705.5 ± 46.5 | 804.5 ± 60.4 | 955.1 ± 35.2 | 1020.6 ± 28.8 | 866.0 ± 38.2 |
| | $k_{cat}/K_m$ ($s^{-1}mM^{-1}$) | *166.23 ± 9.9* | 131.5 ± 11.9 | 100.8 ± 5.8 | 62.2 ± 2.9 | 29.5 ± 1.0 | 23.0 ± 1.8 |
| V169C/E173L/I246A | $K_m$ (mM) | 5.7 ± 0.58 | 5.3 ± 0.6 | 5.9 ± 0.4 | 7.1 ± 0.7 | 14.2 ± 1.1 | 19.2 ± 1.4 |
| | $k_{cat}$ ($s^{-1}$) | 748.7 ± 58.4 | 776.4 ± 33.2 | 805.7 ± 0.9 | 860.9 ± 28.1 | 963.6 ± 32.6 | 937.9 ± 46.0 |
| | $k_{cat}/K_m$ ($s^{-1}mM^{-1}$) | 131.1 ± 3.07 | 146.1 ± 11.4 | 136.9 ± 9.6 | 121.1 ± 8.4 | 68.1 ± 2.9 | 49.0 ± 1.5 |

Table S2. The rationale for the generation of B8CYA8 mutants. The location of the mutations and the intended effects on hydrophobicity and side-chain size are listed.

| **Mutants** | **Location of mutation** | **Effect on Hydrophobicity** | **Effect on side-chain size** |
|---|---|---|---|
| W122F | Glycone region | Increase | Similar |
| V169C | Aglycone region | Decrease | Similar |
| W168A | | Decrease | Decrease |
| W168R | | Decrease | Similar |
| E173L | | Increase | Similar |
| E173A | | Increase | Decrease |
| H180F | Gatekeeper region | Increase | Increase |
| H180K | | Decrease | Similar |
| I246A | | Decrease | Decrease |
| A410F | | Increase | Increase |
| A410K | | Decrease | Increase |

Table S3. The half-life ($t_{1/2}$) of B8CYA8 and mutants in the absence of glucose (Glc) and residual specific activity in the presence of 1 M Glc (see “Materials and methods” for details). The specific activity of each enzyme variant was measured after a 24 h incubation of the enzyme in glucose at 70 °C. The enzyme specific activity was normalized to the specific activity in the absence of glucose to generate the % Residual activity. The equation for linear decay was used to calculate $t_{1/2}$.

| Mutants | Half-life (min) Without Glc | Residual activity (%) (24 h, 70 °C, 1M Glc) |
|---|---|---|
| WT | 75 | 39.8 ± 7.5 |
| W122F | 330 | 72.2 ± 18 |
| W168A | 38 | 40.9 ± 20 |
| W168R | 64 | 62.8 ± 7.0 |
| V169C | 696 | 4.5 ± 0.6 |
| E173L | 32 | 33.1 ± 6.6 |
| E173A | 89 | 55.2 ± 9.7 |
| H180F | 73 | 40.5 ± 3.3 |
| H180K | 154 | 60.1 ± 8.0 |
| I246A | 300 | 61.4 ± 18 |
| V169C/I246A | 348 | 48.8 ± 5.4 |
| A410F | 93 | 67.6 ± 4.2 |
| A410K | 277 | 71.6 ± 2.3 |
| V169C/E173L | 818 | 54.6 ± 4.4 |
| V169C/E173L/I246A | 462 | 75.3 ± 5.1 |

Table S4. Melting temperature ($T_m$) of mutants measured with and without the presence of glucose (Glc). The $T_m$ of mutants was measured by differential scanning fluorimetry (DSF) using a method previously reported [2]. Errors shown here are standard deviations of three independent reactions. The previously published data [3] are indicated by *italics*.

| **Mutants** | **$T_m$** | | |
|---|---|---|---|
| | **No Glc** | **0.5 M Glc** | **1.0 M Glc** |
| WT | *74.7 ± 0.2* | 80.3 ± 0.1 | 82.0 ± 0.1 |
| W122F | 75.3 ± 0.2 | 79.2 ± 0.1 | 80.7 ± 0.4 |
| W168A | 75.3 ± 0.2 | 79.8 ± 0.1 | 81.1 ± 0.1 |
| W168R | 75.2 ± 0.2 | 79.6 ± 0.2 | 81.4 ± 0.2 |
| V169C | *75.6 ± 0.1* | 80.4 ± 0.2 | 82.2 ± 0.1 |
| E173L | 74.2 ± 0.3 | 79.6 ± 0.3 | 81.2 ± 0.2 |
| E173A | 72.0 ± 0.1 | 77.2 ± 0.3 | 79.2 ± 0.2 |
| H180F | 72.4 ± 0.2 | 76.8 ± 0.3 | 78.8 ± 0.2 |
| H180K | 73.8 ± 0.6 | 78.7 ± 0.1 | 80.5 ± 0.2 |
| I246A | *74.5 ± 0.1* | 79.9 ± 0.3 | 81.6 ± 0.1 |
| E354Q (inactive mutant) | 76.5 ± 0.2 | 79.6 ± 0.4 | 82.1 ± 0.4 |
| A410F | 75.1 ± 1.2 | 79.0 ± 0.4 | 82.5 ± 0.3 |
| A410K | 75.2 ± 0.1 | 80.1 ± 0.1 | 81.9 ± 0.1 |
| V169C/E173L | 74.5 ± 0.2 | 80.8 ± 0.3 | 83.3 ± 2.4 |
| V169C/I246A | *75.0 ± 0.1* | 77.9 ± 0.1 | 79.6 ± 0.3 |
| V169C/E173L/I246A | 75.1 ± 0.1 | 80.2 ± 0.6 | 82.3 ± 0.2 |

Table S5. Glucose generated by synergistic action of B8CYA8 (3 µg per assay) with commercial cellulase cocktail (20 µg per assay) from *Trichoderma viride* (Sigma, St. Louis, USA), The glucose was generated through the hydrolysis of Avicel 5 % (w/v) at 37 °C and pH 5.0 and the glucose quantitated by GOD-POD assay. β-glucosidase from sweet almond (3 µg per assay) was used as a control.

| Sample | Glc (mM) |
|---|---|
| Cellulase | 3.7 ± 0.3 |
| + wild-type | 4.4 ± 0.1 |
| + V169C | 5.5 ± 0.3 |
| + V169C/E173L | 6.1 ± 0.4 |
| + V169C/I246A | 6.2 ± 0.2 |
| + V169C/E173L/I246A | 6.8 ± 0.1 |
| + Almond β-glucosidase | 4.1 ± 0.2 |

Table S6. Cooperativity of B8CYA8 and mutants in the absence and presence of 0.5 M and 1.5 M glucose. The kinetic measurements of B8CYA8 and its mutants were done in the presence of glucose and the hill coefficient value generated by non-linear regression fit of allosteric sigmoidal cooperativity in GraphPad Prism. Error bars show mean standard error in the mean.

| **Enzyme** | **Hill coefficient** | | |
|---|---|---|---|
| | **0 Glc** | **0.5 M Glc** | **1.5 M Glc** |
| Wild type | 1.3 ± 0.2 | 1.3 ± 0.0 | 1.3 ± 0.0 |
| W122F | 0.5 ± 0.1 | 1.1 ± 0.1 | 1.1 ± 0.1 |
| W168A | 1.6 ± 0.3 | 1.2 ± 0.1 | 1.1 ± 0.1 |
| W168R | 1.2 ± 0.1 | 1.2 ± 0.2 | 1.2 ± 0.1 |
| E173A | 0.8 ± 0.1 | 0.9 ± 0.1 | 1.4 ± 0.1 |
| E173L | 0.8 ± 0.1 | 1.2 ± 0.2 | 1.4 ± 0.1 |
| V169C | 1.6 ± 0.2 | 1.2 ± 0.1 | 1.3 ± 0.1 |
| H180K | 0.4 ± 0.0 | 1.0 ± 0.3 | 1.1 ± 0.1 |
| H180F | 1.2 ± 0.1 | 1.5 ± 0.3 | 1.3 ± 0.0 |
| I246A | 1.4 ± 0.2 | 1.3 ± 0.1 | 1.1 ± 0.0 |
| A410K | 1.3 ± 0.1 | 1.5 ± 0.2 | 1.2 ± 0.1 |
| A410F | 0.6 ± 0.1 | 1.4 ± 0.3 | 1.4 ± 0.1 |

Table S7. List of primers used for site-directed mutations of B8CYA8

| S. no. | Primers | Sequence |
|---|---|---|
| 1 | W122F reverse | 5' CGCTTGCGGCAAATCGAAATGATATAAAGTGATC 3' |
| 2 | W168A reverse | 5’ GTGACTCATAACGAACCGGCGGTTGTGGCTTTTGAAGG 3’ |
| 3 | W168R reverse | 5’ TGACTCATAACGAACCGAGGGTTGTGGCTTTTGAA 3’ |
| 4 | V169C reverse | 5’ GACCTTCAAAAGCCACACACCACGGTTCGTTATG3’ |
| 5 | E173A reverse | 5’ CCGAATGCATGACCTGCAAAAGCCACAACC 3’ |
| 6 | E173L reverse | 5' GCCGAATGCATGACCGAGAAAAGCCACAACCCACG 3' |
| 7 | V169C/E173L reverse | 5’CGAATGCATGACCTAAAAAAGCCACACCACGGTTCG 3’ |
| 8 | H180K reverse | 5’ TTTTGTACCAGGGGCTTTATTGCCGAATGCATG 3’ |
| 9 | H180F reverse | 5’TTTTGTACCAGGGGCAAAATTGCCGAATGCATG 3’ |
| 10 | I246A reverse | 5’ CAGAAACCATGCATTAGCGTAGTCATCCAGCAATG3’ |
| 11 | E354Q forward | 5’ CCTTTATACATTACTCAGAACGGTGCCGCG 3’ |
| 12 | A410F forward | 5’ GATAACTTTGAATGGAAATATGGTTACTCGAAG 3’ |
| 13 | A410K forward | 5’ GATAACTTTGAATGGTTTTATGGTTACTCGAAG 3’ |

Figure S1. The ligand-binding surface-displayed with docked glucose in the active site pocket of B8CYA8. The hydrogen atoms linked to C1, C3, C5, and C6 carbon of glucose make close contact with amino acid residues at the binding pocket. The figure was prepared using Pymol [4].

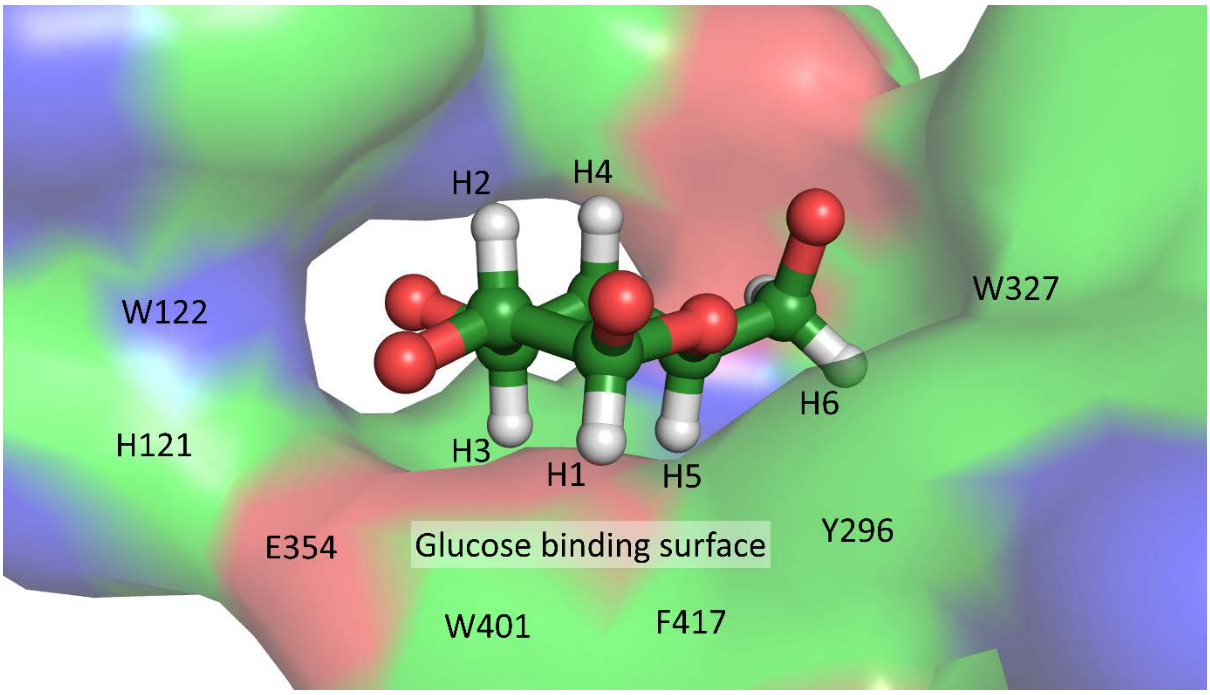

Figure S2. Michaelis-Menten plot of B8CYA8 (wild-type) in the presence of glucose and sucrose. a: B8CYA8 in the presence of 0 M (black), 0.25 M (red), 0.5 M (orange), 0.75 M (green), and 1 M (blue) glucose indicate an increase in $K_m$ and $k_{cat}$ ($V_{max}$) with increasing concentration of glucose. b: In the presence of similar concentrations of sucrose, $K_m$ remains almost unchanged, but $k_{cat}$ ($V_{max}$) increases. This increase is less than at similar concentrations of glucose.

| Concentration (M) | $K_m$ (mM) | | $k_{cat}$ ( $s^{-1}$) | |
|---|---|---|---|---|
| | Glucose | Sucrose | Glucose | Sucrose |
| 0 | 1.8 ± 0.1 | | 339 ± 28 | |
| 0.25 | 04.4 ± 0.6 | 1.7 ± 0.1 | 482 ± 28 | 349 ± 08 |
| 0.5 | 08.1 ± 0.3 | 2.0 ± 0.1 | 572 ± 05 | 399 ± 19 |
| 0.75 | 16.0 ± 1.3 | 2.2 ± 0.1 | 676 ± 18 | 429 ± 05 |
| 1.0 | 23.4 ± 2.2 | 2.0 ± 0.0 | 753 ± 23 | 361 ± 25 |

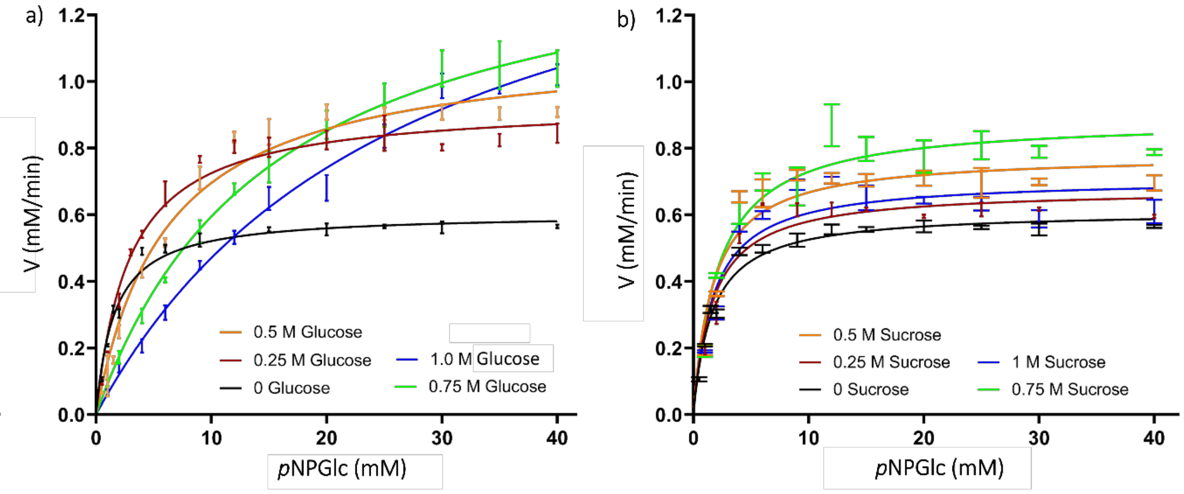

Figure S3. Thin layer chromatography (TLC) of the B8CYA8 catalysed hydrolysis products of *p*NPGlc in the presence of glucose. The reaction products were analyzed on a pre-activated silica plates using a solvent mixture of a n-butanol, n-propanol, ethanol and water (2:3:3:2). Lane 1 represents the carbohydrate standards- *p*NPGlc, glucose (Glc), cellobiose (Clb) and Lactose (Lac). Lane 2-8 hydrolysis product of B8CYA8 with *p*NPGlc (20 mM) as substrate in the presence of 2: Glc 100 mM,3: Glc 250 mM, 4: Glc 500 mM, 5: Glc 750 mM, 6: Glc 1000 mM, 7: 1400 mM and 8: Lactose 100 mM.

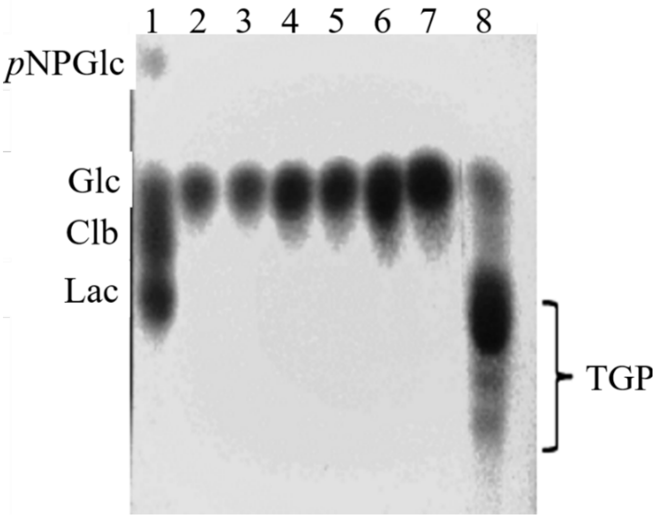

Figure S4. The main parts of the active site pocket including the catalytic residues are shown through a surface representation of the cross-section of the active site pocket of B8CYA8 with its catalytic residues and binding subsites. The PDB coordinates (4PTX) was used [5].

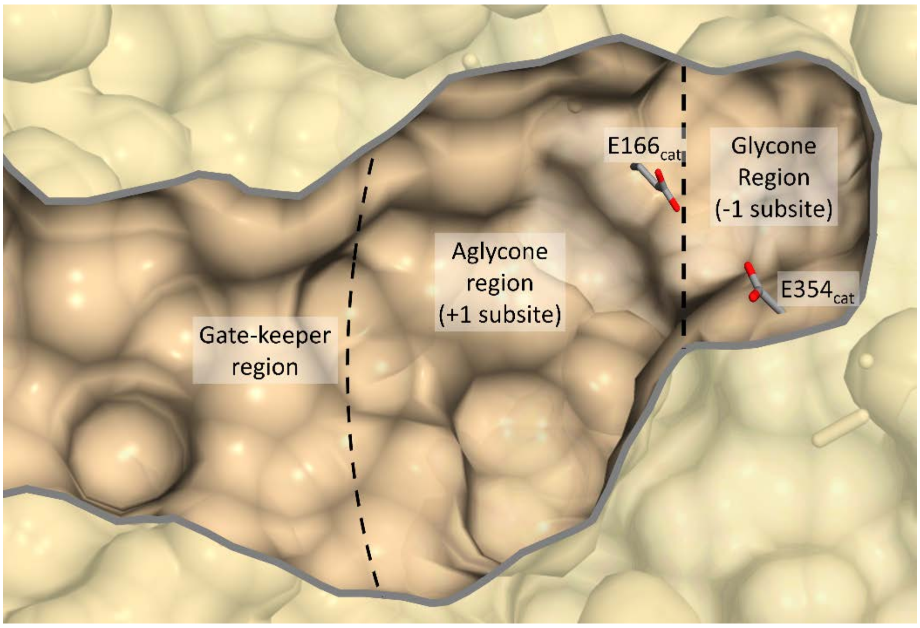

Figure S5. Specific activity of the B8CYA8 gatekeeper mutants on the substrate *p*NPGlc in the presence of a different concentration of glucose under optimum assay condition of each mutant and appropriate substrate concentrations for each. The reaction rate of mutants in the absence of glucose was taken as 100.

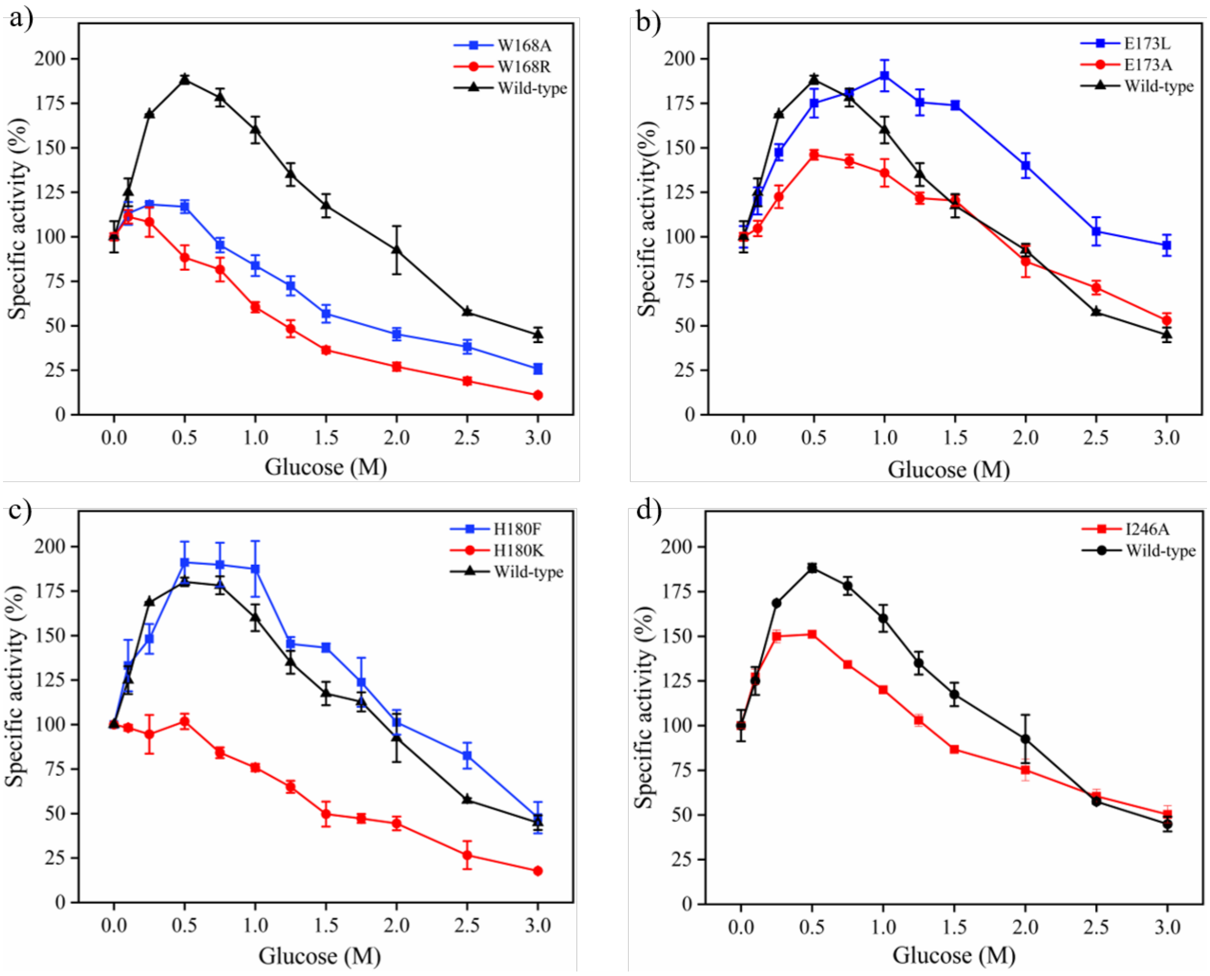
